# Harnessing the Anti-Cancer Natural Product Nimbolide for Targeted Protein Degradation

**DOI:** 10.1101/436998

**Authors:** Jessica N. Spradlin, Xirui Hu, Carl C. Ward, Scott M. Brittain, Michael D. Jones, Lisha Ou, Milton To, Andrew Proudfoot, Elizabeth Ornelas, Mikias Woldegiorgis, James A. Olzmann, Dirksen E. Bussiere, Jason R. Thomas, John A. Tallarico, Jeffrey M. McKenna, Markus Schirle, Thomas J. Maimone, Daniel K. Nomura

## Abstract

Nimbolide, a terpenoid natural product derived from the Neem tree, impairs cancer pathogenicity across many types of human cancers; however, the direct targets and mechanisms by which nimbolide exerts its effects are poorly understood. Here, we used activity-based protein profiling (ABPP) chemoproteomic platforms to discover that nimbolide reacts with a novel functional cysteine crucial for substrate recognition in the E3 ubiquitin ligase RNF114. Nimbolide impairs breast cancer cell proliferation in-part by disrupting RNF114 substrate recognition, leading to inhibition of ubiquitination and degradation of the tumor-suppressors such as p21, resulting in their rapid stabilization. We further demonstrate that nimbolide can be harnessed to recruit RNF114 as an E3 ligase in targeted protein degradation applications and show that synthetically simpler scaffolds are also capable of accessing this unique reactive site. Our study highlights the utility of ABPP platforms in uncovering unique druggable modalities accessed by natural products for cancer therapy and targeted protein degradation applications.

## Introduction

Natural products from organisms such as plants and microbes are a rich source of therapeutic lead compounds^1–5^. The characterization of their biological activities has resulted in myriad medications for disparate pathologies such as cancer, bacterial and fungal infections, inflammation, and diabetes^1–5^. Among natural products there exists a subset of covalently acting molecules that bear electrophilic moieties capable of undergoing essentially irreversible reactions with nucleophilic amino acid hotspots within proteins to exert their therapeutic activity. Examples of these natural products include penicillin, which irreversibly inhibits serine transpeptidases inducing anti-bacterial activity, or wortmannin, which covalently modifies a functional lysine on PI3-kinase to inhibit its activity^5–7^. Discovering druggable hotspots targeted by anti-cancer and covalently-acting natural products can not only yield new cancer drugs and therapeutic targets but can also reveal unique insights into modalities accessed by natural products in protein targets that are often considered undruggable or difficult to tackle with standard drug discovery efforts. One example of a druggable modality that would be difficult to predict *a priori* is FK506 or Tacrolimus that inhibits peptidylprolyl isomerase activity by binding to FKBP12 thus creating a FKBP12-FK506 complex that modulates T cell signaling via inhibition of calcineurin^8^. Gaining insights into nature’s strategies for engaging protein targets can thus provide access to new perspectives on what may be considered druggable.

In this study, we investigated the mechanism of action of the natural product nimbolide, a limonoid natural product derived from the Neem tree (*Azadirachta indica*) (**Figure 1A**)^9^. Nimbolide has been shown to exert multiple therapeutic effects and possesses a cyclic enone capable of reacting with cysteines^10–12^. In the context of cancer, nimbolide has been shown to inhibit tumorigenesis and metastasis without causing any toxicity or unwanted side effects across a wide range of cancers^10, 11, 13–17^. While previous studies suggest that nimbolide impairs cancer pathogenicity through modulating signaling pathways and transcription factors linked to survival, growth, invasion, angiogenesis, inflammation, and oxidative stress the direct targets of nimbolide still remain poorly characterized^10, 15, 18–22^.

**Figure 1.**
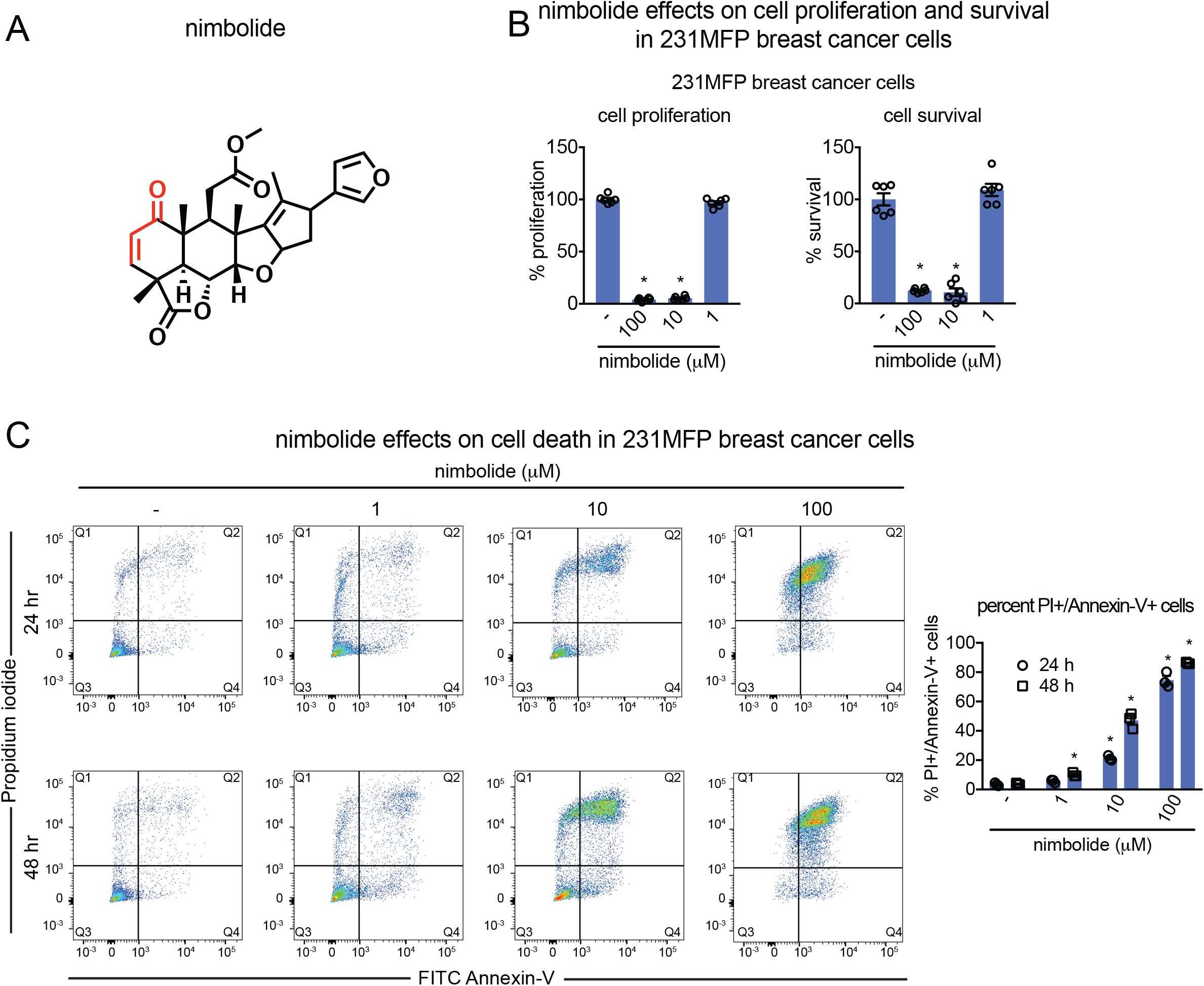
Nimbolide impairs breast cancer cell proliferation or survival. **(A)** Structure of nimbolide. Nimbolide possesses a cyclic enone that is potentially cysteine-reactive. **(B)** 231MFP breast cancer cell proliferation in serum-containing media and serum-free cell survival. **(C)** Percent of propidium iodide and Annexin-V-positive (PI+/Annexin-V+) cells assessed by flow cytometry after treating 231MFP cells with DMSO vehicle or nimbolide for 24 or 48 h. Shown on the left panels are representative FACS data. On the right bar graph are noted late-stage apoptotic cells defined as defined as FITC+/PI+ cells. Data shown in **(A** and **B)** are average ± sem, n=3-6 biological replicates/group. Statistical significance was calculated with unpaired two-tailed Student’s t-tests. Significance is expressed as *p<0.05 compared to vehicle-treated controls.

Identifying direct protein targets of complex natural products remains challenging and often requires synthesizing analogs of these compounds bearing photoaffinity and enrichment handles, a task which is synthetically challenging and has the potential to alter the activity of the molecule^23–25^. Even with the generation of such probes, identifying the specific amino acid site targeted by natural products is challenging. In this study, we utilized activity-based protein profiling (ABPP) chemoproteomic approaches to map the proteome-wide targets of nimbolide in breast cancer cell proteomes. Using ABPP platforms, we reveal that one of the primary targets of nimbolide in breast cancer cells is cysteine-8 (C8) of the E3 ubiquitin ligase RNF114. Covalent modification of RNF114 by nimbolide leads to impaired ubiquitination and degradation of its substrate—the tumor suppressor p21—leading to its rapid stabilization. Intriguingly, we show that this apparent inhibition of RNF114 activity is through impaired substrate recognition giving rise to the possibility that nimbolide could be used as an E3 ligase recruitment module for targeted protein degradation. Strategies for chemically induced degradation of targets of interest in cells is rapidly gaining interest in drug discovery, including the development of bifunctional molecules referred to as “proteolysis-targeting chimeras” (PROTACs) or “heterobifunctional degraders” that consist of a protein-targeting ligand linked to an E3 ligase recruiter to bring an E3 ligase to a protein of interest to ubiquitinate and mark the target for degradation in a proteasome-dependent manner^26–34^. We demonstrate that nimbolide can be used to recruit RNF114 to other protein substrates for targeted protein degradation applications. Using chemoproteomics-enabled covalent ligand screening platforms, we also identify more synthetically tractable compounds that can similarly react with C8 of RNF114 and phenocopy nimbolide action.

## Results

### Effects of nimbolide on breast cancer cell survival, proliferation, and apoptosis

Though nimbolide has been shown to impair cancer pathogenicity across many different types of human cancers, we chose to focus on elucidating the role of nimbolide in triple-negative breast cancers (TNBC). TNBCs are devoid of estrogen, progesterone, and HER2 receptors and are amongst the most aggressive cancers with the worst clinical prognosis^35, 36^. Very few targeted therapies currently exist for TNBC patients. Uncovering new therapeutic modalities in TNBCs would thus potentially contribute significantly to reducing breast cancer-associated mortalities. Consistent with previous reports showing anti-cancer activity in breast cancer cells, nimbolide impaired cell proliferation or serum-free cell survival in 231MFP and HCC38 TNBC cells (**Figure 1B**, **Figure S1**)^14, 15, 17^. We also show that these effects on 231MFP and HCC38 viability are due to a significant increase in apoptotic cells with nimbolide treatment, assessed by flow cytometry analysis (**Figure 1C**, **Figure S1**).

### Using ABPP platforms to map druggable hotspots targeted by nimbolide in breast cancer cells

To interrogate the mechanisms by which nimbolide impairs breast cancer pathogenicity, we applied a chemoproteomic platform termed isotopic tandem orthogonal proteolysis-enabled activity-based protein profiling (isoTOP-ABPP) to determine the specific protein targets of nimbolide. Pioneered by Cravatt and Weerapana, isoTOP-ABPP uses reactivity-based chemical probes to map reactive, functional, and ligandable hotspots directly in complex proteomes^37–40^. When used in a competitive manner, covalently-acting small-molecules can be competed against binding of broad reactivity-based probes to facilitate the rapid discovery of both proteins and ligandable sites targeted by the covalently-acting compound^40–47^ (**Figure 2A**). In this study, we treated breast cancer cells *in situ* with vehicle or nimbolide followed by competitive labeling of proteomes with a cysteine-reactive alkyne-functionalized iodoacetamide probe (IA-alkyne), after which isotopically light or heavy cleavable enrichment handles were appended to probe-labeled proteins for isoTOP-ABPP analysis. Probe-modified tryptic peptides were analyzed by liquid chromatography-mass spectrometry (LC-MS/MS) and light to heavy tryptic probe-modified peptide ratios, representing control versus treated IA-alkyne labeled sites, were quantified. IsoTOP-ABPP analysis of ligandable hotspots targeted by *in situ* nimbolide treatment in 231MFP TNBC cells showed one primary target showing an isotopic ratios >4 that was significantly engaged by nimbolide—the E3 ubiquitin ligase RNF114 (**Figure 2B; Table S1)**. Importantly, RNF114 knockdown using three independent small interfering RNA (siRNA) resembled the anti-proliferative effects of nimbolide in 231MFP cells (**Figure 2C-2D, Figure S2**). Further demonstrating that RNF114 contributes to the anti-proliferative effects of nimbolide, RNF114 knockdown led to significant resistance to nimbolide-mediated anti-proliferative effects (**Figure 2E**, **Figure S2**). While nimbolide likely possesses many additional targets beyond RNF114 that are not accessible with isoTOP-ABPP approaches, our results suggested that RNF114 was a novel target of nimbolide and that RNF114 was in-part responsible for the anti-proliferative effects of this natural product. We thus chose to focus further characterization efforts on the interactions between nimbolide and RNF114.

**Figure 2.**
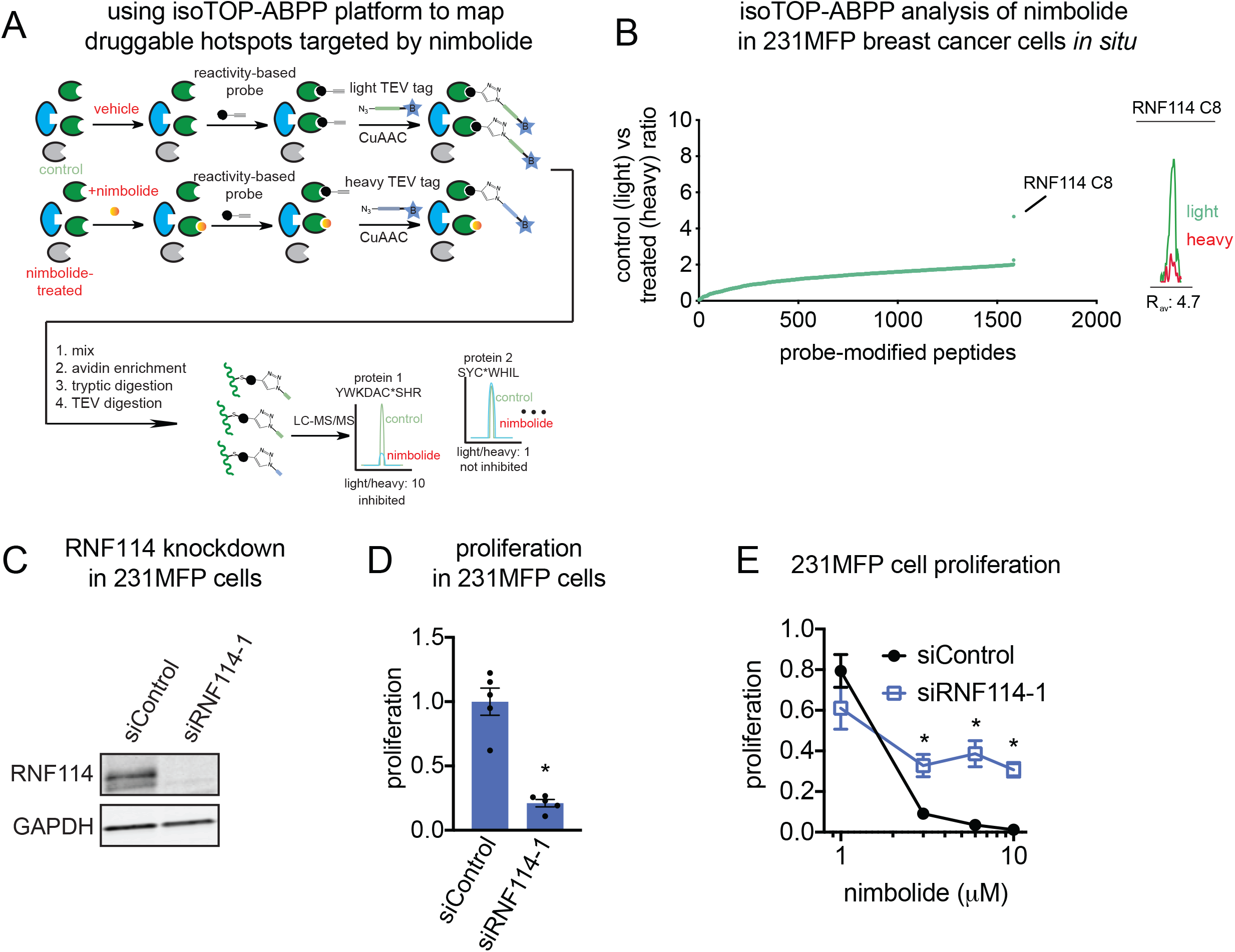
isoTOP-ABPP analysis of nimbolide in 231MFP breast cancer cell proteomes reveal RNF114 as a target. **(A)** Schematic of isoTOP-ABPP in which 231MFP breast cancer cells were treated with DMSO or nimbolide (10 μM, 1.5 h *in situ*), after which cells were harvested and proteomes were labeled *ex situ* with IA-alkyne (100 μM, 1 h), followed by appendage of isotopically light (for DMSO-treated) or heavy (for nimbolide-treated) TEV protease cleavable biotin-azide tags by copper-catalyzed azide-alkyne cycloaddition (CuAAC). Control and treated proteomes were subsequently combined in a 1:1 ratio, probe-labeled proteins were avidin-enriched, digested with trypsin, and probe-modified tryptic peptides were eluted by TEV protease, analyzed by LC-MS/MS, and light to heavy probe-modified peptide ratios were quantified. **(B)** isoTOP-ABPP analysis of nimbolide (10 μM) in 231MFP breast cancer cells *in situ* analyzed as described in **(A).** Light vs heavy isotopic probe-modified peptide ratios are shown in the left plot where the top average ratio is 4.7 corresponding to C8 of RNF114. Shown on the right is a representative MS1 light vs heavy peak for the probe-modified peptide bearing C8 of RNF114. **(C)** RNF114 knockdown by short interfering RNA (siRNA) targeting RNF114 validated by Western blotting of RNF114 compared to siControl 231MFP cells. GAPDH expression is shown as a loading control. Shown gel is a representative gel from n=3 biological replicates/group. **(D)** 231MFP cell proliferation after 24 h in siControl and siRNF114 cells assessed by Hoechst stain. **(E)** Nimbolide effects on 231MFP siControl and siRNF114 231MFP breast cancer cells. Nimbolide effects on 231MFP siControl and siRNF114 231MFP breast cancer cells. Cells were treated with DMSO vehicle or nimbolide for 24 h after which proliferation was assessed by Hoechst stain. Data for siControl or siRNF114 group was normalized to the respective DMSO vehicle control in each group. Data shown in **(D-E)** are average ± sem. Data shown in **(B-E)** are from n=3-5 biological replicates/group. Statistical significance was calculated with unpaired two-tailed Student’s t-tests. Significance in **(D)** is expressed as *p<0.05 compared to siControl cells. Significance in **(E)** is expressed as *p<0.05 compared to the corresponding nimbolide treatment concentration from siControl groups.

### Biochemical characterization of nimbolide interactions with RNF114

RNF114 is an E3 ubiquitin ligase of the RING family^48, 49^. The site on RNF114 identified by isoTOP-ABPP as the target of nimbolide, C8, falls within the N-terminal region of the protein, predicted to be intrinsically disordered, and resides outside of the two annotated zinc finger domains (**Figure 3A**). Intrigued by the apparent targeting of an intrinsically disordered region of a protein by a natural product, we sought to investigate the interaction between nimbolide and RNF114.

**Figure 3.**
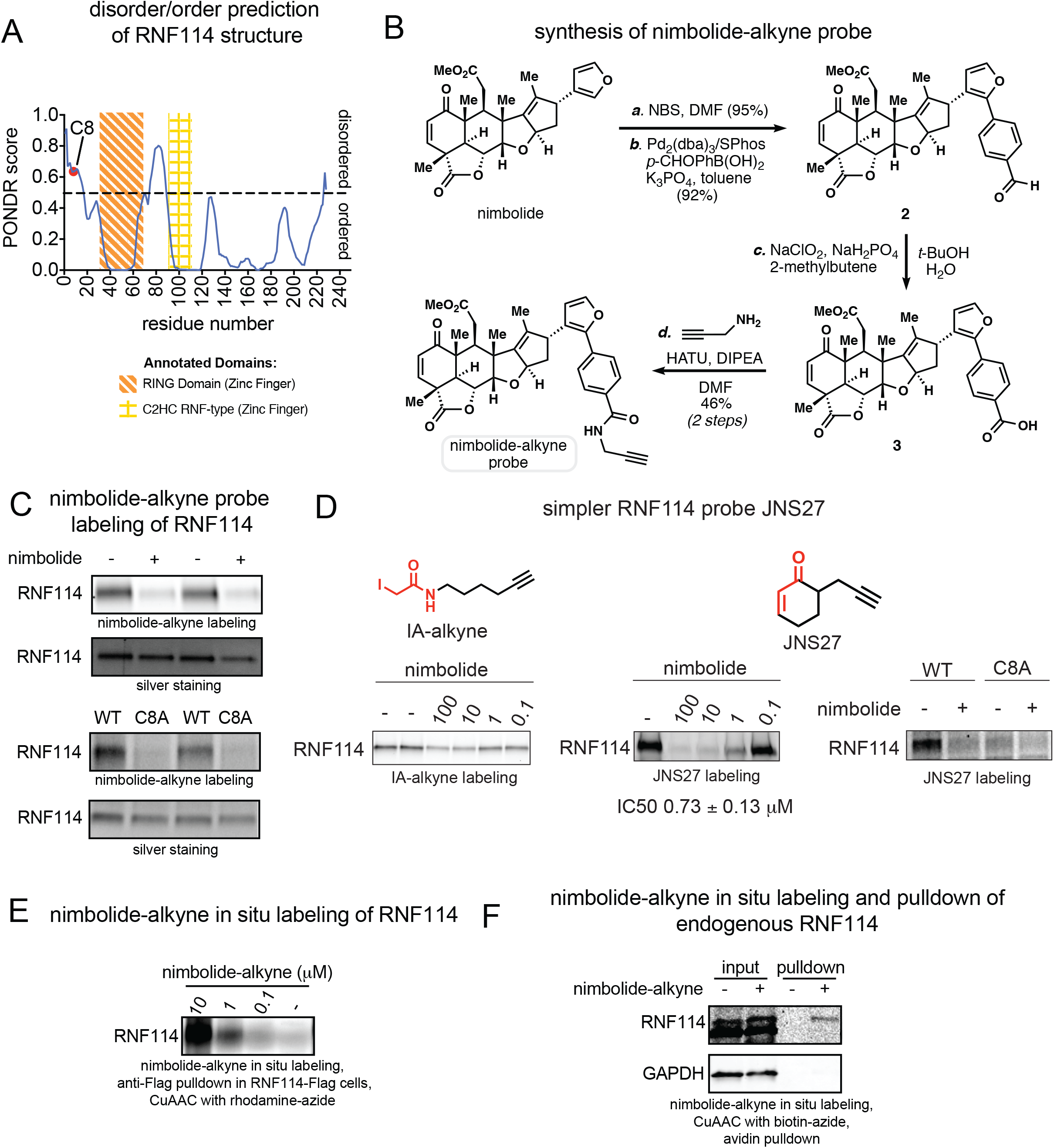
Nimbolide reacts covalently with C8 of RNF114. **(A)** Nimbolide targets an intrinsically disordered region within RNF114 as assessed by PONDR. **(B)** Route for synthesis of the alkyne-functionalized nimbolide probe. **(C)** Gel-based ABPP analysis of pure human RNF114 protein labeled with nimbolide probe. In the upper two panels, pure RNF114 protein was pre-incubated with DMSO vehicle or nimbolide (100 μM, 30 min) prior to labeling with the nimbolide probe (10 μM, 1 h) in PBS. In the lower two panels, pure wild-type and C8A mutant RNF114 protein were labeled with the nimbolide probe (10 μM, 1 h) in PBS with 1 mg/ml BSA. For both experiments, shown are nimbolide-alkyne labeling and silver staining of RNF114. **(D)** Gel-based ABPP analysis of nimbolide competition against IA-alkyne (10 μM) or JNS27 (50 μM) labeling of pure RNF114 protein. Structures of IA-alkyne and JNS27 probes are shown with reactive moieties highlighted in red. Shown also is gel-based ABPP analysis of nimbolide (50 μM) competition against JNS27 labeling of wild-type and C8A mutant RNF114 protein. In these experiments, DMSO or nimbolide was pre-incubated for 30 min prior to probe labeling for 1 h. **(E)** Nimbolide-alkyne labeling of Flag-RNF114 in 231MFP cells. 231MFP cells stably expressing a Flag-tagged RNF114 were treated with DMSO vehicle or nimbolide-alkyne for 2 h. RNF114 was subsequently enriched from harvested cell lysates and then rhodamine-azide was appended onto probe-labeled proteins by CuAAC, after which nimbolide-alkyne labeling was visualized by SDS/PAGE and in-gel fluorescence. **(F)** Nimbolide-alkyne labeling of endogenous RNF114 in 231MFP cells. 231MFP cells were treated with DMSO vehicle or nimbolide-alkyne (50 μM) for 1.5 h. Biotin-azide was appended to probe-labeled proteins by CuAAC, and these proteins were subsequently avidin-enriched. Resulting pulled down proteins were analyzed by SDS/PAGE and Western blotting for RNF114. Shown in the gel are RNF114 and GADPH expression from input proteome and nimbolide-alkyne labeled protein pulldown. Gels shown in **(C-F)** are representative gels from n=3 biological replicates/group.

Because isoTOP-ABPP is a competitive assay between the covalently-acting molecule and a broader reactive probe, it is an indirect read-out of potential nimbolide-targeting hotspots. To confirm that nimbolide directly targeted RNF114, we synthesized the alkyne-functionalized probe shown in three steps from nimbolide (**Figure 3B**). Selective bromination at C2 of the furan moiety and subsequent Suzuki coupling with 4-Formylphenylboronic acid afforded the aryl-aldehyde **2**, which was converted to its corresponding carboxylic acid (**3**) via Pinnick oxidation. Finally, coupling of **3** and propargyl amine (HATU, DIPEA) afforded the target probe. We confirmed that this nimbolide probe reacts with pure human RNF114 protein as shown by labeling of the protein on a denaturing SDS/PAGE gel. This labeling event was also competed by nimbolide and abrogated in the C8A RNF114 mutant (**Figure 3C**). As the nimbolide-alkyne probe is a synthetically laborious probe that requires the rather expensive nimbolide as a starting material, we also sought to develop a more simple probe that could label C8 of RNF114. In initial studies trying to validate RNF114 interactions with nimbolide, where we competed nimbolide against IA-alkyne labeling of RNF114, full inhibition of labeling was not observed, likely due to alkylation of multiple cysteines by IA-alkyne beyond C8. Thus, we synthesized a more tailored alkyne-functionalized cyclic enone probe JNS27, which showed selective labeling of C8 on RNF114 as evidenced by lack of labeling of C8A RNF114 mutant protein (**Figure 3D**). With JNS27, we were able to demonstrate full competition of JNS27 labeling with nimbolide with a 50 % inhibitory concentration (IC50) of 0.73 μM (**Figure 3D**). To demonstrate that nimbolide interactions with RNF114 are not completely driven by reactivity, but rather also through additional interactions with the protein, we show that nimbolide competes against labeling of RNF114 with a rhodamine-functionalized iodoacetamide probe (IA-rhodamine) at far lower concentrations compared to the simpler JNS27 probe or iodoacetamide (**Figure S2**) with an IC50 of 0.55, 22, and >100 μM, respectively (**Figure S2**). Furthermore, a direct mass adduct of nimbolide was detected on the tryptic peptide containing C8 on RNF114 by LC-MS/MS after incubation of pure protein with the natural product (**Figure S2**). All of these experiments taken together provide evidence that nimbolide modifies an intrinsically disordered region of RNF114 at C8 through a covalent bond formation.

We next performed two complementary experiments to demonstrate that nimbolide directly engaged RNF114 in 231MFP breast cancer cells. First, we showed dose-responsive nimbolide-alkyne labeling of RNF114 *in situ* by treating Flag-tagged RNF114 expressing cells with the probe, followed by enrichment of Flag-tagged RNF114 and appending rhodamine-azide onto probe-labeled protein by CuAAC and visualizing by gel-based ABPP methods (**Figure 3E**, **Figure S2**). We observed robust *in situ* nimbolide-alkyne labeling with 10 μM of probe, but also observed lower but significant probe labeling even down to 100 nM (**Figure 3E**, **Figure S2**). Second, we also showed *in situ* labeling of endogenous RNF114 with the nimbolide-alkyne probe by treating cells with the probe, followed by appending biotin-azide onto probe-labeled proteins by CuAAC and visualizing RNF114 pulldown by Western blotting (**Figure 3F**). In this case, significant pulldown was observed at 50 μM of the nimbolide-alkyne probe but not lower concentrations, likely reflecting the need to saturate RNF114 labeling in order to detect RNF114 using this method. In this experiment, we show that an unrelated protein such as GAPDH is not enriched by the nimbolide-alkyne probe (**Figure 3F**). Using these latter conditions, we also performed a complementary quantitative proteomic profiling study to identify any additional proteins that may be enriched from *in situ* labeling of 231MFP cells with the nimbolide-alkyne probe **(Table S2)**. Due to acute cytotoxicity issues, we were not able perform *in situ* competition studies with higher concentrations of nimbolide itself. In this study, we showed that RNF114 was one of the proteins enriched by the nimbolide-alkyne probe by >10-fold compared to no-probe controls. We also identified 114 additional proteins that were enriched by the probe by >10-fold compared to no-probe controls. These proteins represent additional potential covalent or non-covalent targets that may contribute to the anti-proliferative effects of nimbolide. These enriched targets may also represent proteins with low to partial degrees of engagement with nimbolide, whereas isoTOP-ABPP experiments are meant to identify higher engagement targets. Additionally, these proteins may include probe-specific targets or proteins that may be enriched indirectly through interactions with direct nimbolide-labeled proteins **(Table S2)**. Nonetheless, we show that RNF114 is engaged by the nimbolide-alkyne probe in breast cancer cells, even down to 100 nM, and that the anti-proliferative effects of nimbolide are at least in-part mediated by RNF114.

### Effects of nimbolide on RNF114 activity and substrate binding *in vitro* and *in situ*

RNF114 has been previously shown to ubiquitinate and degrade the tumor suppressor p21, among other substrates^48, 50, 51^. In an *in vitro* reconstituted system, nimbolide inhibited both RNF114 autoubiquitination and p21 ubiquitination activity (**Figure 4A**). The RNF114 C8A mutation did not affect basal RNF114 autoubiquitination activity, but attenuated the inhibition observed with nimbolide treatment (**Figure 4B**). Previous characterization of RNF114 suggested that the N-terminus may be involved in substrate recognition^48^. Consistent with this premise, we found that the amount of p21 co-immunoprecipitated with RNF114 was reduced by nimbolide, suggesting that the apparent inhibition of RNF114 may be due to impaired substrate recognition, rather than inhibition of its catalytic activity (**Figure 4C**). We further demonstrated that nimbolide treatment in 231MFP cells stabilized p21 protein expression within 1 hour in a dose-responsive manner, with no significant changes to p53 levels (**Figure 4D**, **Figure S3**). The elevated levels of CDKN1A observed were not due to transcriptional downregulation of mRNA levels, as p21 mRNA levels remained unchanged with nimbolide treatment (**Figure S3**). To identify other potential substrates of RNF114, we also performed a tandem mass tagging (TMT)-based quantitative proteomic experiment on 231MFP cells treated with nimbolide. Consistent with our Western blotting data, we observed CDKN1A (p21) as one of the proteins that were significantly elevated >2-fold with nimbolide treatment (**Figure 4E, Table S3**). We also observed the levels of several other targets that were heightened by nimbolide treatment, including CDKN1C (p57), PEG10, and CTGF. Beyond CDKN1A (p21) and CDKN1C (p57) which have previously been reported as potential RNF114 substrates^48^, we conjectured that PEG10 and CTGF may also represent additional novel substrates of RNF114 (**Figure 4E, Table S3**). Consistent with this premise, we demonstrated in an *in vitro* RNF114 ubiquitination assay that RNF114 ubiquitinates PEG10 and CTGF and that this ubiquitination is inhibited by nimbolide (**Figure 4F**). As both p21 (CDKN1A) and p57 (CDKN1C) are tumor suppressors that, when elevated, promote cell cycle arrest and apoptosis^52, 53^, we postulated that the stabilization of both of these tumor suppressors may be responsible for the anti-proliferative effect of nimbolide. We demonstrated that dual knockdown of p21 (CDKN1A) and p57 (CDKN1C) results in significant attenuation in nimbolide-mediated anti-proliferative effects in 231MFP breast cancer cells (**Figure 4G-4H).** Collectively, these data suggest that nimbolide reacts with an intrinsically disordered C8 of RNF114 in breast cancer cells to disrupt RNF114-substrate recognition, leading to inhibition of ubiquitination of its substrates such as CDKN1A and CDKN1C, leading to their stabilization, and impaired cell proliferation in breast cancer cells.

**Figure 4.**
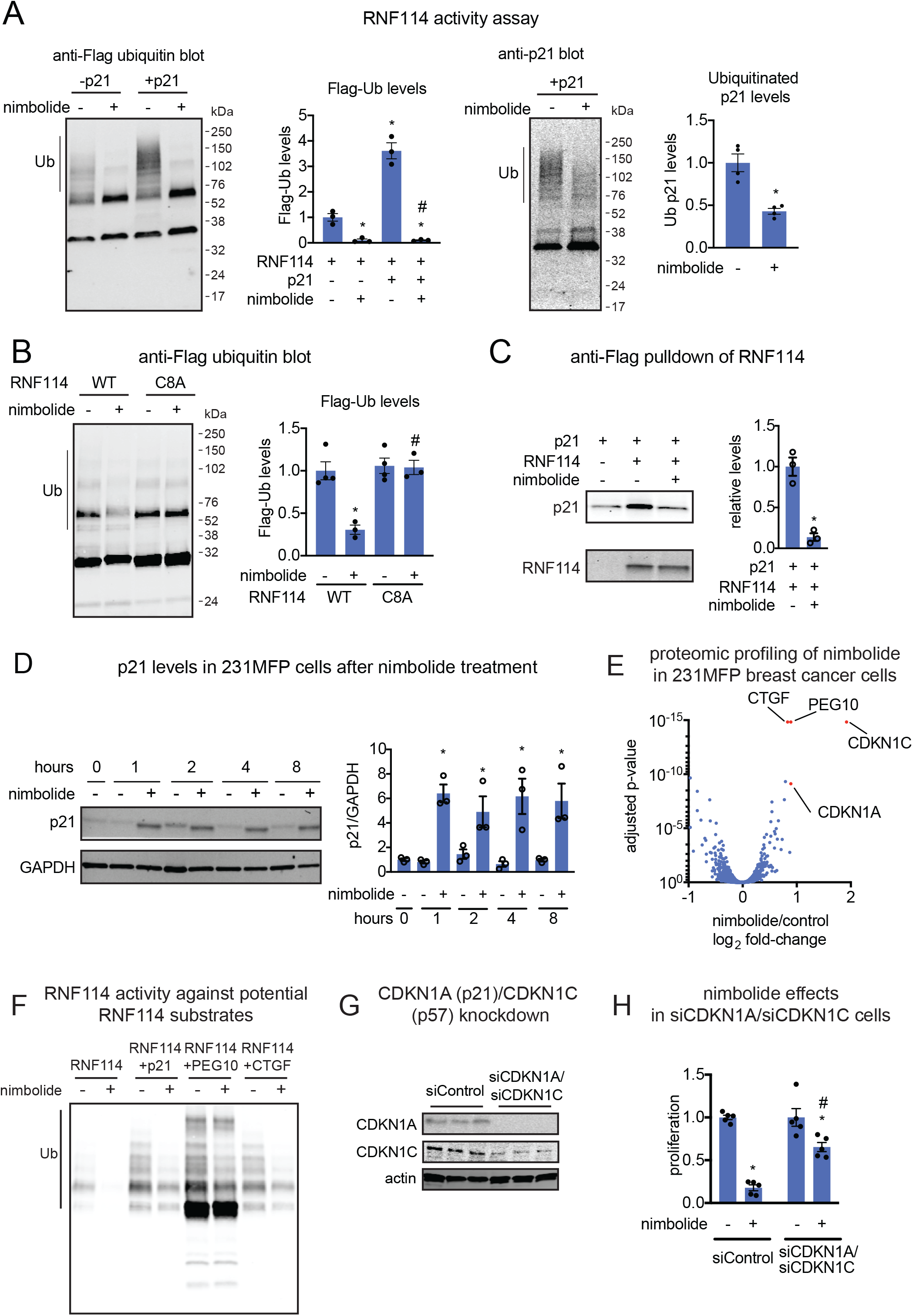
Nimbolide inhibits RNF114 activity through disrupting substrate recognition. **(A)** RNF114 ubiquitination assay with pure GST-Ube1, GST-UBE2D1, and RNF114 protein, Flag-Ubiquitin and ATP with or without addition of p21 and blotting against Flag-ubiquitin (left) or p21 (right). DMSO or Nimbolide (100 μM) was pre-incubated with RNF114, before the addition of the E1 and E2 enzymes, Flag-ubiquitin and ATP to start the reaction. Anti-Flag-ubiquitin and Ubiquitinated p21 are quantified in bar graphs by densitometry. **(B)** RNF114 autoubiquitination assay with DMSO or nimbolide (100 μM) treatment with wild-type or C8A mutant RNF114. Anti-Flag ubiquitin are quantified in bar graphs by densitometry. **(C)** In an *in vitro* incubation of pure RNF114 and p21 protein, Flag-RNF114 pulldown and p21 enrichment were inhibited by nimbolide (100 μM). **(D)** 231MFP cells were treated with nimbolide (100 μM). Shown are p21 levels in DMSO control or nimbolide-treated cells. **(E)** Tandem mass tag (TMT)-based quantitative proteomic profiling of 231MFP breast cancer cells treated with DMSO vehicle or nimbolide (100 nM) for 12 h. Shown in red are proteins that are significantly heightened in levels >2-fold. **(F)** RNF114 ubiquitination assay with pure GST-Ube1, GST-UBE2D1, and RNF114 protein, Flag-Ubiquitin and ATP with the addition of p21 (CDKN1A), PEG10, or CTGF and blotting against Flag-ubiquitin. DMSO or Nimbolide (100 μM) was pre-incubated with RNF114, before the addition of the E1 and E2 enzymes, Flag-ubiquitin and ATP to start the reaction. **(G)** p21 (CDKN1A) and p57 (CDKN1C) expression in siControl and siCDKN1A/siCDKN1C 231MFP cells, assessed by Western blotting, alongside actin as a loading control. Data shown in **(G)** are for 6397 proteins quantified with 2 or more unique peptides in triplicate treatments, see **Table S3** for details. **(H)** 231MFP cell proliferation in siControl or siCDKN1A/siCDKN1C cells treated with DMSO vehicle or nimbolide (6 μM) for 24 h. Gels shown in **(A-D,** and **F)** are representative images from n=3 biological replicates/group. Data shown in **(A-D,** and **H)** are average ± sem, n=3-5 biological replicates/group. Statistical significance was calculated with unpaired two-tailed Student’s t-tests. Significance is expressed as *p<0.05 compared to RNF114 DMSO vehicle treated control group for Flag-Ub blot and compared to RNF114/p21 DMSO vehicle control group for **(A)**, compared to WT DMSO vehicle-treated control in **(B)**, compared to the p21/RNF114 group in **(C)**, against vehicle-treated control groups for each time point in **(D),** and compared to vehicle-treated controls for each group in **(H)**. Significance expressed as #p<0.05 compared to RNF114/p21 vehicle-treated control group in **(A)**, compared to WT nimbolide-treated group in **(B)**, and compared to nimbolide-treated siControl cells in **(H)**.

### Using nimbolide as an RNF114-recruiter for targeted protein degradation

Since nimbolide targets a potential substrate recognition site, we conjectured that nimbolide could be used to recruit RNF114 to other protein substrates for proteasomal degradation through the development of heterobifunctional degraders using nimbolide as an RNF114 recruiter. To demonstrate feasibility, two degraders formed by linking nimbolide to the Bromodomain and extra-terminal (BET) family inhibitor JQ1 were synthesized (**Figure 5A**; **Figure S4**). Prior studies have demonstrated efficient proteasome-dependent degradation of BET family members and in particular BRD 4 with JQ1-based degraders linked to either a cereblon-recruiter thalidomide or a VHL recruiter^27, 30^. Previously prepared acid **3** was coupled to JQ1-functionalized amines containing both longer (PEG-based) and shorter (alkyl-based) spacer units, arriving at degraders **XH1** and **XH2 (Figure 5A**; **Figure S4**). We show that **XH2** still binds to RNF114 with an IC50 of 0.24 μM (**Figure 5B**). While **XH1** did not show appreciable BRD4 degradation, **XH2** treatment in 231MFP cells led to BRD4 degradation after a 12 h (**Figure 5C**; **Figure S4**). Interestingly, **XH2** showed less degradation at 1 μM compared to 0.1 and 0.01 μM and MZ1 showed less degradation at 20 μM compared to 10, 1, and 0.1 μM, which we attribute to the “hook effect” previously reported with other degraders including the previously reported JQ1-based degrader MZ1 that utilizes a VHL recruiter^26, 27^. To confirm proteasome-dependence of BRD4 degradation, we showed that the **XH2**-mediated degradation of BRD4 was attenuated by pre-treatment of cells with the proteasome inhibitor bortezomib (BTZ) (**Figure 5D**). **XH2**-mediated BRD4 degradation was also prevented by pre-treatment with an E1 ubiquitin-activating enzyme inhibitor (TAK-243) or pre-competing with the BRD4 inhibitor JQ1 (**Figure 5E**, **Figure S4**). However, treatment with a translation inhibitor (emetine) had no effect of the observed degradation of BRD4 (**Figure S4**). To further demonstrate that the degradation of BRD4 by **XH2** was through the specific recruitment of RNF114, we showed that degradation of BRD4 by **XH2**, but not MZ1, was not observed in RNF114 knockout (KO) HAP1 cells compared to wild-type (WT) HAP1 cell counterparts (**Figure 5F**). We further showed that re-expression of wild-type RNF114 in HAP1 RNF114 knockout cells led to the restoration of BRD4 degradation by XH2 (**Figure S4**). The selectivity of **XH2**-mediated degradation of proteins was demonstrated using TMT-based quantitative proteomic profiling experiment to assess changes in protein expression. We showed that **XH2** selectively degrades BRD4 in 231MFP breast cancer cells while sparing the other identified BET family members BRD 2 and 3 (**Figure 5G; Table S3**). Of note, we also observed several proteins that showed increased levels upon **XH2** treatment including CDKN1A, CDKN1C, PEG10, and CTGF, which were observed as elevated in levels with nimbolide treatment alone. There were also additional proteins that were upregulated, such as DSG1, FN1, FLG2, and CASP14. These upregulated proteins may be potential substrates of RNF114 with stabilization stemming from the nimbolide portion of XH2, as is likely the case with CDKN1A, CDKN1C, PEG10 and CTGF. The other targets may be downstream transcriptional effects stemming from JQ1-mediated BRD4 inhibition and degradation. Our results indicate that nimbolide reactivity with RNF114 can be exploited to recruit this E3 ligase to other protein substrates, such as BRD4, to ubiquitinate and selectively degrade them.

**Figure 5.**
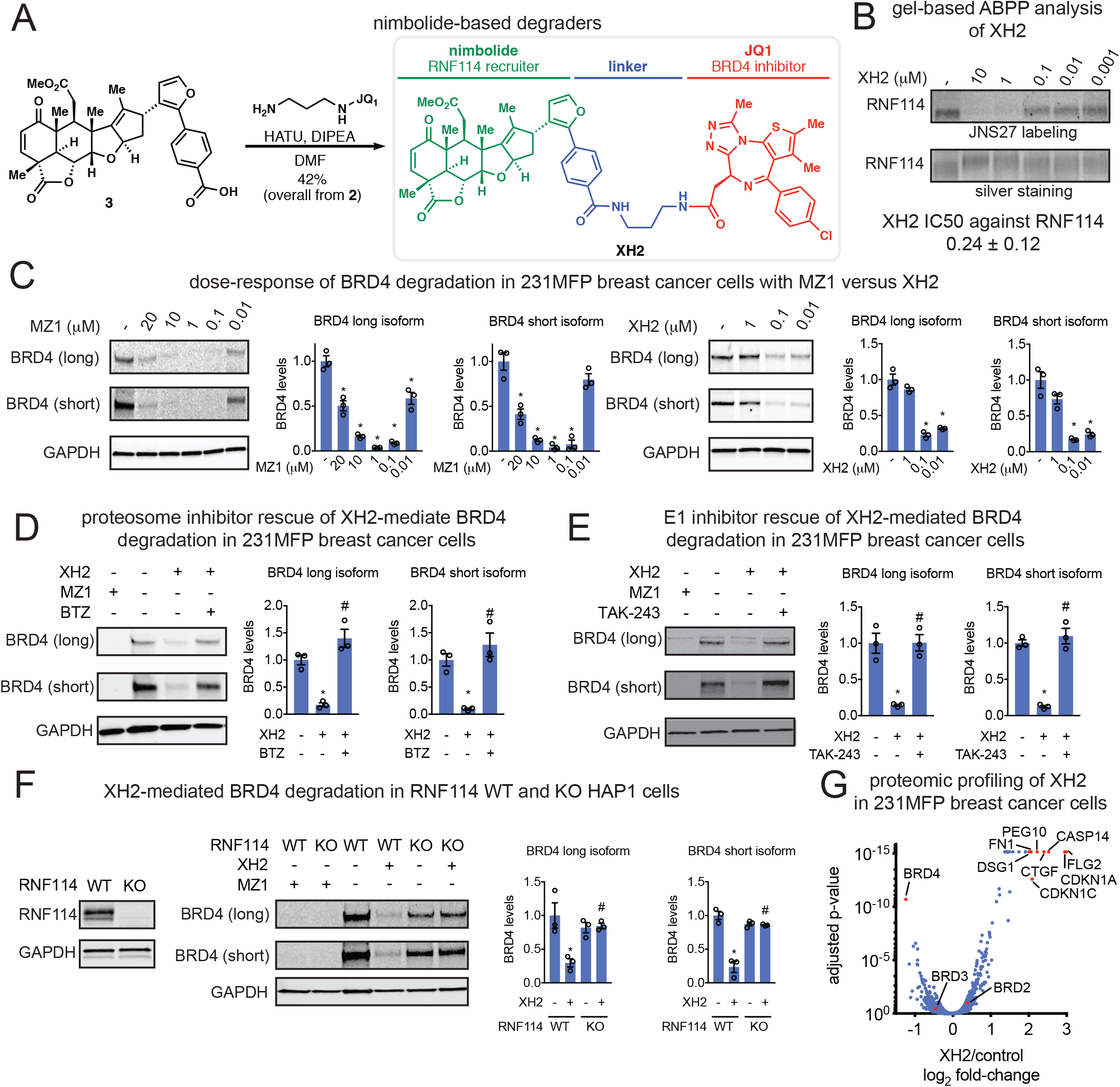
Nimbolide can be used to recruit RNF114 for targeted protein degradation of BRD4. **(A)** Route for synthesizing XH2, a nimbolide-based degrader consisting of nimbolide as an RNF114 recruiter, a linker, and the BRD4 inhibitor JQ1. **(B)** Gel-based ABPP analysis of XH2 against RNF114. RNF114 was pre-incubated with DMSO vehicle or XH2 for 30 min prior to JNS27 labeling (50 μM) for 1 h followed by appending rhodamine-azide by CuAAC, SDS/PAGE, and analysis of in-gel fluorescence. IC50 value noted below. **(C)** BRD4 levels in 231MFP breast cancer cells treated with MZ1 versus XH2 treatment for 12 h. **(D-E)** BRD4 levels in 231MFP breast cancer cells pre-treated with DMSO vehicle or proteasome inhibitor bortezomib (BTZ) (1 μM) **(D)** or E1 ubiquitin activating enzyme inhibitor TAK-243 (10 μM) **(E)** 30 min prior to and also during treatment with MZ1 (1 μM) or XH2 (100 nM) treatment (100 nM) for 12 h. **(F)** RNF114 and BRD4 expression in RNF114 wild-type (WT) or knockout (KO) HAP1 cells treated with DMSO vehicle, MZ1 (1 μM), or XH2 (100 nM) for 12 h. **(G)** Tandem mass tag (TMT)-based quantitative proteomic profiling of 231MFP breast cancer cells treated with DMSO vehicle or XH2 (100 nM) for 12 h. Long and short BRD4 isoforms in **(C-F)** were visualized by SDS/PAGE and Western blotting, quantified by densitometry, and normalized to GAPDH loading controls in **(C-F)**. Gels shown in **(C-F)** are representative images from n=3 biological/group. Data shown in **(G)** are for 5797 proteins quantified with 2 or more unique peptides in triplicate treatments, see **Table S3** for details. Data shown in **(C-F)** are average ± sem. Statistical significance in **(C-F)** was calculated with unpaired two-tailed Student’s t-tests. Significance in **(C-F)** is expressed as *p<0.05 compared to the vehicle-treated control groups and #p<0.05 compared to XH2-treated groups in **(C-F)** or to the WT XH2-treated group **(F)**. Statistical significance in **(G**) are described in the methods section and p-values are reported in **Table S3.**

### Chemoproteomics-enabled covalent ligand screening to identifying covalent ligands against RNF114

With the insight gained that C8 of RNF114 has potential to be exploited for cancer therapy and targeted protein degradation applications, we searched for more synthetically tractable covalent ligands that similarly target RNF114. To achieve this goal, we screened a library of cysteine-reactive covalent ligands against RNF114 using a moderate-throughput gel-based ABPP approach, in which covalent ligands are competed against JNS27 labeling of RNF114, followed by appending a rhodamine-azide and analysis by SDS/PAGE and in-gel fluorescence. Of the approximately 200 cysteine-reactive covalent ligands screened against RNF114, the acrylamide **EN62** emerged as a promising hit (**Figure S5**, **Figure S6, Table S4**). **EN62** inhibited RNF114 autoubiquitination activity in a C8-dependent manner (**Figure S6**). While **EN62** is an early hit compound that requires substantial improvements to cell permeability, potency, and selectivity, this covalent ligand represents a starting point for more synthetically tractable covalent ligand scaffolds that target C8 of RNF114 for targeted protein degradation applications.

## Discussion

Collectively, we show compelling evidence that nimbolide impairs breast cancer pathogenicity in-part through targeting a substrate recognition domain at C8 within RNF114 to inhibit p21 ubiquitination and degradation, leading to its stabilization. We demonstrate that nimbolide targeting of C8 on RNF114 can be used to recruit RNF114 for targeted protein degradation applications and degradation of BRD4 with a nimbolide-JQ1 degrader **XH2** is possible.

We report here that nimbolide disrupts RNF114 interactions with one of its endogenous substrate p21 and we also show that p21 levels are rapidly stabilized in breast cancer cells in a p53-independent manner. Several other E3 ligases have also been reported to degrade p21, including SCF^Skp2^, CRL4^Cdt2^, and CHIP under varying conditions during cell cycle or exogenous stress^54–56^. Previous studies have shown that RNF114 expression is elevated at late G1 phase to regulate p21 and p57 levels and is crucial for G1-to-S phase transition^48^. Other RNF114 substrates that have been reported include TAB1 involved in maternal-to-zygotic transition and A20 involved in NF-κB activity and T cell activation^57, 58^. In cancer contexts or other cell and tissue types, nimbolide may thus have additional activities through regulating the levels of other RNF114 protein substrates. Intriguingly, we show CTGF and PEG10 may represent additional substrates of RNF114. Further studies are required to establish these additional proteins as direct and endogenous substrates of RNF114. Furthermore, while we show that RNF114 is one target of nimbolide that at least in-part mediates its effects upon proliferation and targeted protein degradation, we also show with the nimbolide-alkyne probe that nimbolide likely possesses many additional protein targets. These additional targets may include other covalent interactions with cysteines that are not labeled by the IA-alkyne cysteine-reactive probe in isoTOP-ABPP experiments, may represent covalent interactions with other amino acids, may represent reversible interactions with additional protein targets, or may include proteins are indirectly enriched from interactions with direct nimbolide-labeled proteins. Despite nimbolide possessing multiple potential targets, we demonstrate that RNF114 is an important and functional target of nimbolide in breast cancer cells that can be exploited for targeted protein degradation applications. We also convincingly show that the BRD4 degradation observed with the nimbolide-based degrader is driven through RNF114.

Our results also demonstrate that nimbolide functionally targets an intrinsically disordered region within RNF114. Solving the structure of RNF114 covalently modified with nimbolide has thus far proven challenging, but future studies investigating whether nimbolide induces order in the N-terminus would provide insights into the ligandability of intrinsically disordered and undruggable protein targets and strategies for potentially targeting other E3 ligases.

Targeted protein degradation has emerged as a formidable and effective drug discovery paradigm for targeting proteins for elimination through the proteasome^26, 28^. One of the challenges, however, is that there are only a small number of E3 ligase recruiters that have been developed among the approximately 600 E3 ligases in the human genome^59^. These E3 ligase recruiters include the thalidomide-type immunomodulatory drugs (IMiD) that recruit cereblon, ligands that recruit VHL, nutlins that recruit MDM2, and ligands that recruit cIAP^26, 28^. Here, we report that nimbolide can be used as a novel RNF114 recruiter for targeted protein degradation applications. It should be possible to optimize the performance of this degrader class *via* further linker modifications, an area of the molecule already determined to be important. It may also be possible to utilize more synthetically tractable covalent ligands capable of targeting C8 of RNF114, such as EN62, as RNF114 recruiters. Since nimbolide targets a substrate recognition domain within RNF114, it will also be of future interest to determine whether nimbolide may act as a molecular glue to recruit and degrade neo-substrates, as has been reported for the IMiDs^60–62^.

Interestingly, we did not observe degradation of BRD2 and 3 under conditions that led to significant reduction of BRD4 levels despite the high homology in their respective BET bromodomains. Various levels of selectivity within the BET family members and the structural basis thereof have been reported for JQ1-based degraders with Cereblon and VHL-recruiting modules^27, 30, 63, 64^. While a more detailed investigation of the selectivity of XH2 and its structural basis is outside the scope of this study, it serves as another example how availability of additional E3-moieties for a given substrate recognition modules can aid in tuning efficacy and selectivity of degraders targeting a given protein of interest.

Overall, our study further demonstrates the utility of using ABPP-based chemoproteomic platforms to identify unique druggable modalities exploited by natural products. Intriguingly, we show that a natural product can functionally access an E3 ligase protein-protein interaction site for potential cancer therapy and targeted protein degradation applications and remarkably does so in an intrinsically disordered region of the protein. Our study also showcases how covalent ligand screening approaches can be utilized to identify more synthetically tractable small-molecules that act similarly to more complex natural products and that covalent ligands may be able to access other E3 ligases to expand the scope of E3 ligase recruiters.

## Methods

### Cell Culture

The 231MFP cells were obtained from Prof. Benjamin Cravatt and were generated from explanted tumor xenografts of MDA-MB-231 cells as previously described^65^. HCC38 and HEK293T cells were obtained from the American Type Culture Collection. HEK293T cells were cultured in DMEM containing 10% (v/v) fetal bovine serum (FBS) and maintained at 37°C with 5% CO_2_. 231MFP were cultured in L15 medium containing 10% FBS and maintained at 37°C with 0% CO_2_. HCC38 cells were cultured in RPMI medium containing 10% FBS and maintained at 37°C with 5% CO_2_. HAP1 RNF114 wild-type and knockout cell lines were purchased from Horizon Discovery. The RNF114 knockout cell line was generated by CRISPR/Cas9 to contain a frameshift mutation in a coding exon of RNF114. HAP1 cells were grown in Iscove’s Modified Dulbecco’s Medium (IMDM) in the presence of 10 % FBS and penicillin/streptomycin.

### Survival and Proliferation Assays

Cell survival and proliferation assays were performed as previously described using Hoechst 33342 dye (Invitrogen) according to manufacturer’s protocol and as previously described^66^. 231MFP cells were seeded into 96-well plates (40,000 for survival and 20,000 for proliferation) in a volume of 150 μl and allowed to adhere overnight. Cells were treated with an additional 50 μL of media containing 1:250 dilution of 1000 × compound stock in DMSO. After the appropriate incubation period, media was removed from each well and 100 μl of staining solution containing 10% formalin and Hoechst 33342 dye was added to each well and incubated for 15 min in the dark at room temperature. After incubation, staining solution was removed, and wells were washed with PBS before imaging. Studies with HCC38 cells were also performed as above but were seeded with 20,000 cells for survival and 10,000 cells for proliferation.

### Assessing Apoptosis in Breast Cancer Cells

Apoptotic analyses were performed 24 or 48 h following cell exposure (6 cm plates, 1 × 10^6^ cells) to DMSO vehicle or compound-containing serum-free media using flow cytometry to measure the percentage of early apoptotic cells (Annexin V positive, propidium iodide negative) and late apoptotic cells (Annexin V positive, propidium iodide positive), as described previously^42^. Data analysis was performed using FlowJo software.

### IsoTOP-ABPP Chemoproteomic Studies

IsoTOP-ABPP studies were done as previously reported^37, 40, 44, 67^. Cells were lysed by probe sonication in PBS and protein concentrations were measured by BCA assay^68^. For *in situ* experiments, cells were treated for 90 min with either DMSO vehicle or covalently-acting small molecule (from 1000X DMSO stock) before cell collection and lysis. Proteomes were subsequently labeled with IA-alkyne labeling (100 μM) for 1 h at room temperature. CuAAC was used by sequential addition of tris(2-carboxyethyl)phosphine (1 mM, Sigma), tris[(1-benzyl-1H-1,2,3-triazol-4-yl)methyl]amine (34 μM, Sigma), copper (II) sulfate (1 mM, Sigma), and biotin-linker-azide—the linker functionalized with a TEV protease recognition sequence as well as an isotopically light or heavy valine for treatment of control or treated proteome, respectively. After CuAAC, proteomes were precipitated by centrifugation at 6500 × g, washed in ice-cold methanol, combined in a 1:1 control/treated ratio, washed again, then denatured and resolubilized by heating in 1.2 % SDS/PBS to 80°C for 5 minutes. Insoluble components were precipitated by centrifugation at 6500 × g and soluble proteome was diluted in 5 ml 0.2% SDS/PBS. Labeled proteins were bound to avidin-agarose beads (170 μl resuspended beads/sample, Thermo Pierce) while rotating overnight at 4°C. Bead-linked proteins were enriched by washing three times each in PBS and water, then resuspended in 6 M urea/PBS (Sigma) and reduced in TCEP (1 mM, Sigma), alkylated with iodoacetamide (IA) (18 mM, Sigma), then washed and resuspended in 2 M urea and trypsinized overnight with 0.5 μg/μl sequencing grade trypsin (Promega). Tryptic peptides were eluted off. Beads were washed three times each in PBS and water, washed in TEV buffer solution (water, TEV buffer, 100 μM dithiothreitol) and resuspended in buffer with Ac-TEV protease and incubated overnight. Peptides were diluted in water and acidified with formic acid (1.2 M, Spectrum) and prepared for analysis.

### Mass Spectrometry Analysis

Peptides from all chemoproteomic experiments were pressure-loaded onto a 250 μm inner diameter fused silica capillary tubing packed with 4 cm of Aqua C18 reverse-phase resin (Phenomenex # 04A-4299) which was previously equilibrated on an Agilent 600 series HPLC using gradient from 100% buffer A to 100% buffer B over 10 min, followed by a 5 min wash with 100% buffer B and a 5 min wash with 100% buffer A. The samples were then attached using a MicroTee PEEK 360 μm fitting (Thermo Fisher Scientific #p-888) to a 13 cm laser pulled column packed with 10 cm Aqua C18 reverse-phase resin and 3 cm of strong-cation exchange resin for isoTOP-ABPP studies. Samples were analyzed using an Q Exactive Plus mass spectrometer (Thermo Fisher Scientific) using a 5-step Multidimensional Protein Identification Technology (MudPIT) program, using 0%, 25 %, 50 %, 80 %, and 100 % salt bumps of 500 mM aqueous ammonium acetate and using a gradient of 5-55 % buffer B in buffer A (buffer A: 95:5 water:acetonitrile, 0.1 % formic acid; buffer B 80:20 acetonitrile:water, 0.1 % formic acid). Data was collected in data-dependent acquisition mode with dynamic exclusion enabled (60 s). One full MS (MS1) scan (400-1800 m/z) was followed by 15 MS2 scans (ITMS) of the nth most abundant ions. Heated capillary temperature was set to 200 °C and the nanospray voltage was set to 2.75 kV.

Data was extracted in the form of MS1 and MS2 files using Raw Extractor 1.9.9.2 (Scripps Research Institute) and searched against the Uniprot human database using ProLuCID search methodology in IP2 v.3 (Integrated Proteomics Applications, Inc)^69^. Cysteine residues were searched with a static modification for carboxyaminomethylation (+57.02146) and up to two differential modifications for methionine oxidation and either the light or heavy TEV tags (+464.28596 or +470.29977, respectively). Peptides were required to be fully tryptic peptides and to contain the TEV modification. ProLUCID data was filtered through DTASelect to achieve a peptide false-positive rate below 5%. Only those probe-modified peptides that were evident across two out of three biological replicates were interpreted for their isotopic light to heavy ratios. For those probe-modified peptides that showed ratios >2, we only interpreted those targets that were present across all three biological replicates, were statistically significant, and showed good quality MS1 peak shapes across all biological replicates. Light versus heavy isotopic probe-modified peptide ratios are calculated by taking the mean of the ratios of each replicate paired light vs. heavy precursor abundance for all peptide spectral matches (PSM) associated with a peptide. The paired abundances were also used to calculate a paired sample t-test p-value in an effort to estimate constancy within paired abundances and significance in change between treatment and control. P-values were corrected using the Benjamini/Hochberg method.

### Knockdown of RNF114, p21 and p57 by RNA Interference (RNAi) in 231MFP Cells

RNAi was performed by using siRNA purchased from Dharmacon specific to RNF114 or p21 and p57 for dual knockdown. In brief, 231MFP cells were seeded at a density of 5×10^4^ cells per mL full media in a 96-well format. On day 0, 231MFP cells were transfected with corresponding small interfering RNA (siRNA) vs. non-targeting negative control (Dharmacon, ON-TARGETplus Non-targeting Pool D-001810-10-05) duplexes at 50 nM using Dharmafect 1 (Dharmacon) as a transfection reagent for 48 h. Thereafter (day 2), transfection media was supplemented with fresh DMSO or compound containing full L15 media and cultured for an additional 24 to 48 h before undergoing cell viability testing as described above. At the time of treatment (day 2), RNA was extracted for analysis by qRT-PCR and lysates were obtained for Western blotting confirmation of knockdown.

Targeting Sequences:

siRNF114-1: GUGUGAAGGCCACCAUUAA (Dharmacon J-007024-08-0002)
siRNF114-2: GCUUAGAGGUGUACGAGAA (Dharmacon J-007024-05-0002)
siRNF114-3: GCACGGAUACCAAAUCUGU (Dharmacon J-007024-06-0002)
sip21: AGACCAGCAUGACAGAUUU (Dharmacon J-003471-12-0002)
sip57: CUGAGAAGUCGUCGGGCGA (Dharmacon J-003244-13-0002)

### Gene Expression by qPCR

Total RNA was extracted from cells using Trizol (Thermo Fisher Scientific). cDNA was synthesized using MaximaRT (Thermo Fisher Scientific) and gene expression was confirmed by qPCR using the manufacturer’s protocol for Fisher Maxima SYBR Green (Thermo Fisher Scientific) on the CFX Connect Real-Time PCR Detection System (BioRad). Primer sequences for Fisher Maxima SYBR Green were derived from Primer Bank. Sequences of primers are as follows:

RNF114 Forward: AAT GTT CCA AAC CG
RNF114 Reverse: TTG CAG TGT TCC AC
p21 Forward: TGT CCG TCA GAA CCC ATG C
p21 Reverse: AAA GTC GAA GTT CCA TCG CTC
Cyclophilin Forward: CCC ACC GTG TTC TTC GAC ATT
Cyclophilin Reverse: GGA CCC GTA TGC TTT AGG ATG A

### Gene Expression by qPCR

Gene expression was confirmed by qPCR using the manufacturer’s protocol for Fisher Maxima SYBR Green. Primer sequences for Fisher Maxima SYBR Green were derived from Primer Bank. Sequences of primers are as follows:

RNF114 Forward: AAT GTT CCA AAC CG
RNF114 Reverse: TTG CAG TGT TCC AC
Cyclophilin Forward: CCC ACC GTG TTC TTC GAC ATT
Cyclophilin Reverse: GGA CCC GTA TGC TTT AGG ATG A

### Gel-Based ABPP

Gel-based ABPP methods were performed as previously described^43, 66, 67, 70^. Recombinant pure human proteins were purchased from Origene. Pure proteins (0.1 μg) were pre-treated with DMSO vehicle or covalently-acting small molecules for 30 min at room temperature in an incubation volume of 50 μL PBS, and were subsequently treated with JNS-1-27 (50 μM final concentration) for 1 h at room temperature. CuAAC was performed to append rhodamine-azide (1 μM final concentration) onto alkyne probe-labeled proteins. Samples were then diluted with 20 μL of 4 × reducing Laemmli SDS sample loading buffer (Alfa Aesar) and heated at 90 °C for 5 min. The samples were separated on precast 4-20% TGX gels (Bio-Rad Laboratories, Inc.). Prior to scanning by ChemiDoc MP (Bio-Rad Laboratories, Inc), gels were fixed in a solution of 10% acetic acid, 30% ethanol for 2 hrs. Inhibition of target labeling was assessed by densitometry using ImageJ.

### Synthesis and Characterization of the Nimbolide-Alkyne Probe and Degraders XH1 and XH2

See supporting information for experimental details.

### Synthesis and Characterization of JNS27

See supporting information for experimental details.

### Covalent Ligand Library

The synthesis and characterization of most of the covalent ligands screened against RNF114 have been previously reported^40, 41, 43, 67^. Synthesis of TRH 1-156, TRH 1-160, TRH 1-167, YP 1-16, YP 1-22, YP 1-26, YP 1-31, YP 1-44 have been previously reported^71–78^. Compounds starting with “EN” were purchased from Enamine LLC. The synthesis and characterization of other covalent ligands not previously reported are described in **Supporting Information.**

### Western Blotting

Antibodies to RNF114 (Millipore Sigma, HPA021184), p21 (Cell Signaling Technology, 12D1), GAPDH (Proteintech Group Inc., 60004-1-Ig), BRD4 (Abcam plc, Ab128874), DYKDDDDK Tag (Cell Signaling Technology, D6W5B) and beta-actin (Proteintech Group Inc., 6609-1-Ig) were obtained from various commercial sources and dilutions were prepared per recommended manufacturers’ procedures. Proteins were resolved by SDS/PAGE and transferred to nitrocellulose membranes using the iBlot system (Invitrogen). Blots were blocked with 5 % BSA in Tris-buffered saline containing Tween 20 (TBST) solution for 1 h at room temperature, washed in TBST, and probed with primary antibody diluted in recommended diluent per manufacturer overnight at 4 °C. Following washes with TBST, the blots were incubated in the dark with secondary antibodies purchased from Ly-Cor and used at 1:10,000 dilution in 5 % BSA in TBST at room temperature. Blots were visualized using an Odyssey Li-Cor scanner after additional washes. If additional primary antibody incubations were required the membrane was stripped using ReBlot Plus Strong Antibody Stripping Solution (EMD Millipore, 2504), washed and blocked again before being reincubated with primary antibody.

### Expression and Purification of Wild-type and C8A Mutant RNF114 Protein

RNF114 was expressed and purified using several methods. In each case, RNF114 activity and sensitivity to nimbolide was confirmed. For the first method, we purchased wild-type mammalian expression plasmids with C-terminal FLAG tag were purchased from Origene (Origene Technologies Inc., RC209752). The RNF114 C8A mutant was generated with Q5 site-directed mutagenesis kit according to manufacturer’s instructions (New England Biolabs, E0554S). Expression and purification conditions were optimized as reported previously^79^. HEK293T cells were grown to 60% confluency in DMEM (Corning) supplemented with 10 % FBS (Corning) and 2 mM L-glutamine (Life Technologies) and maintained at 37 °C with 5% CO_2_. Immediately prior to transfection, media was replaced with DMEM containing 5 % FBS. Each plate was transfected with 20 μg of overexpression plasmid with 100 μg PEI (Sigma). After 48 h cells were collected in TBS, lysed by sonication, and batch bound with anti-DYKDDDDK resin (GenScript, L00432) for 90 min. Lysate and resin was loaded onto a gravity flow column and washed, followed by elution with 250 ng/μL 3 × FLAG peptide (ApexBio, A6001). Purity and concentration were verified by PAGE, UV/Spectroscopy, and BCA assay.

For the second method, DNA encoding the complete human isoform of RNF114 (Uniprot id: Q9Y508) was codon optimized for expression in *E. coli* and synthesized by Integrated DNA Technologies. Desired constructs were amplified from the complete RNF114 sequence with primers that contained 20 base pairs of homology to a pET24a plasmid (Novagen) that had been modified to contain a His_8_-MBP-TEV sequence between the Nde1 and BamH1 restriction sites. PCR products were analyzed using precast 1% agarose gels (Invitrogen), and those of the correct length were purified using a QIAquick Gel Extraction kit (Qiagen). The purified PCR product was assembled into the linearized vector using Gibson Assembly (NEB Gibson Assembly 2x Master Mix), and transformed into *E. coli* 10G chemically competent cells (Lucigen, Middleton, WI). Colonies resistant to kanamycin (Kan) were grown in LB medium, and the plasmid was isolated via Miniprep (Qiagen) prior to being sequence verified using forward and reverse primers.

100 ng of pET24a His_8_–MBP plasmid containing the desired RNF114 construct was transformed into *E. coli* BL21(DE3) chemically competent cells (NEB product # C2530H). The following day, a single transformed colony was used to inoculate 50 mL of nutrient rich LB medium containing kanamycin (50 μg/mL) and was incubated overnight at 37 ℃, with agitation at 250 rpm. The following morning, 1 L of Terrific Broth (TB) supplemented with 50 mM 3-(N-morpholino)propanesulfonic acid (MOPS) pH 7.5, 1 mM zinc chloride, and kanamycin (50 µg/mL) was inoculated with the overnight starter culture to a starting optical density at 600 nm (OD_600_) of 0.1. Cells were grown at 37 ℃ with agitation at 250 rpm until an OD_600_ of 0.8 was achieved. At this stage, expression of the RNF114 fusion protein was induced with 1 mM isopropyl β-D-1-thiogalactopyranoside (IPTG), and the temperature of the incubator was reduced to 18 ℃. Cells were left growing at 18 °C for 18 hours and were subsequently harvested by centrifugation, washed with 1x phosphate buffered saline (PBS) buffer and stored at −20 °C.

10 g of *E. coli* cells containing the overexpressed RNF114 fusion protein were re-suspended in 80 mL of lysis buffer (50 mM Tris, pH 7.5, 150 mM NaCl, 2 Roche protease inhibitor tablets [without EDTA], 200 mM ZnCl_2_, 1 mM DTT) and sonicated on ice, with a cycling time of 60 seconds on, 60 seconds off, over a total sonication time of three minutes. The cell lysate was centrifuged at 18,000 rpm for 20 minutes and the soluble protein was incubated with agitation for four hours at 4 ℃ with 2 mL of Ni-NTA resin, which had been pre-equilibrated with wash buffer (50 mM Tris, pH 7.5, 150 mM NaCl, 200 mM ZnCl_2_, 1 mM DTT, 25 mM imidazole). The cell lysate / Ni-NTA resin mixture was placed into a disposable column and all non-tagged soluble protein, which do not bind to the resin, was collected for a second round of purification. The resin was washed with 25 mL of wash buffer before the His-MBP-RNF114 protein was eluted from the Ni-NTA resin with 25 mL of elution buffer (50 mM tris, pH 7.5, 500 mM imidazole, 200 mM ZnCl_2_, 150 mM NaCl, 1 mM DTT). This process was repeated incubating the collected flow thorough with an additional 2 mL of pre-equilibrated Ni-NTA resin.

His-MBP-RNF114 protein was simultaneously digested at 4 ℃ with TEV protease (100 units / mg MBP-RNF114), and dialyzed overnight against dialysis buffer (50 mM tris, pH 7.5, 150 mM NaCl, 200 mM ZnCl_2_, 1 mM DTT). The following morning two 5 mL His Trap Excel columns assembled in tandem were placed on an Äkta Avant and equilibrated with five column volumes of wash buffer. The TEV cleaved sample was loaded onto the column at a rate of 2 mL/min and the resin was washed with an additional five column volumes of wash buffer. All fractions identified to contain cleaved RNF114 were concentrated to a volume of 5 mL and loaded onto a HiLoad 16/60 Superdex 75 gel filtration column (GE Healthcare) that had been pre-equilibrated with either crystallography buffer, (25 mM Tris, pH 7.5, 137 mM NaCl) or NMR buffer, (20 mM Sodium Phosphate pH 6.8, 150 mM NaCl,). The gel filtration column was run using a flow rate of 0.25 mL/min and 2 mL fractions were collected. Fractions corresponding to peaks eluting from the gel filtration column were analyzed using SDS-PAGE and all fractions containing RNF114 were concentrated to a final concentration of 10 mg/mL, flash frozen in liquid nitrogen and stored at −80 °C until needed.

### Generation of Stably Expressing FLAG-RNF114 231MFP Cell Lines by Lentiviral Transduction

Human RNF114 with C-terminal FLAG tag was inserted into pLenti vector by FastCloning^80^. FLAG-RNF114 lentivirus was generated by co-transfection of FLAG-RNF114, VSV.G and psPAX2 into HEK 293T cells using Lipfectamine 2000 transfection reagent (ThemoFischer). 24h after transfection media was exchanged for DMEM with 10% heat-inactivated serum (HIS) and after an additional 24hr virus-containing media was collected and filtered with 0.45 μM filter onto 231MFP cells with equal volume HIS L15 media, in the presence of 10 μg/mL polybrene (Santa Cruz). 24h after transduction, puromycin (2 μg/mL) was added to cells and stably expressing FLAG-RNF114 were obtained after puromycin selection for 72hr. 231MFP cells stably expressing FLAG-eGFP were generated in parallel as control.

### *In situ* Nimbolide-Alkyne Probe Labeling and FLAG-RNF114 Pulldown

231MFP cells stably expressing FLAG-RNF114 were treated with either vehicle (DMSO) or 100 nM to 10 μM nimbolide alkyne probe for 2-4hr. Cells were harvested in PBS and lysed by sonication. Total protein concentration of lysates were normalized by BCA assay and normalized lysates were incubated 1.5 hr at 4 °C with 50 μL or FLAG-agarose slurry. After incubation samples were transferred to spin columns and washed 3X with 500 μL PBS. Proteins were eluted using 2 50 μL washes of PBS supplemented with 250 ng/μL 3 × FLAG peptide (ApexBio A6001). CuAAC was performed to append rhodamine-azide onto alkyne probe-labeled proteins and after addition of loading buffer samples were separated on precast 4-20% TGX gels (Bio-Rad Laboratories, Inc.) and imaged on ChemiDoc MP (Bio-Rad Laboratories, Inc).

### LC-MS/MS Analysis of RNF114

Purified RNF114 (10 μg) in 50 μL PBS were incubated 30 min at room temperature either with DMSO vehicle or covalently acting compound (100 μM). The DMSO control was then treated with light IA while the compound treated sample was incubated with heavy IA for 1 h each at room temperature (200 μM final concentration, Sigma-Aldrich, 721328). The samples were precipitated by additional of 12.5 µL of 100% (w/v) TCA and the treated and control groups were combined pairwise, before cooling to −80 °C for 1 h. The combined sample was then spun for at max speed for 10 min at 4 °C, supernatant is carefully removed and the sample is washed with ice cold 0.01 M HCl/90 % acetone solution. The pellet was resuspended in 4 M urea containing 0.1 % Protease Max (Promega Corp. V2071) and diluted in 40 mM ammonium bicarbonate buffer. The samples were reduced with 10 mM TCEP at 60 °C for 30 min. The sample was then diluted 50% with PBS before sequencing grade trypsin (1 μg per sample, Promega Corp, V5111) was added for an overnight incubation at 37 °C. The next day the sample was centrifuged at 13200 rpm for 30 min. The supernatant was transferred to a new tube and acidified to a final concentration of 5 % formic acid and stored at −80 °C until MS analysis.

### RNF114 Ubiquitination Assay

Recombinant Myc-Flag-RNF114 proteins were either purified from HEK292T cells as described above or purchased from Origene (Origene Technologies Inc., TP309752). For *in vitro* auto-ubiquitination assay, 0.2 μg of RNF114 in 25 μL of TBS was pre-incubated with DMSO vehicle or the covalently-acting compound for 30 min at room temperature. Subsequently, 0.1 μg of UBE1 (Boston Biochem. Inc, E-305), 0.1 μg UBE2D1 (Boston Bichem. Inc, e2-615), 5 μg of Flag-ubiquitin (Boston Bichem. Inc, u-120) in a total volume of 25 μL Tris buffer containing 2 mM ATP, 10 mM DTT, and 10 mM MgCl_2_ were added to achieve a final volume of 50 μL. For substrate-protein ubiquitination assays, 0.1ug of the appropriate substrate protein, purchased from commercial sources, was added at this stage (p21 and PEG10: Origene, CTGF: R&D Systems). The mixture was incubated at 37 °C with agitation for 1.5 h. 20 μL of Laemmli’s buffer was added to quench the reaction and proteins were analyzed by western blot assay.

### RNF114/p21 Co-Immunoprecipitation

Recombinant Flag-tagged RNF114 was used as bait to precipitate pure recombinant p21 (Origene Technologies Inc., TP309752 and TP720567) using Anti-Flag agarose beads (GenScript Biotech Corp., L00432). One microgram of Flag-RNF114 was added to 50 μL of TBS, followed by the addition of nimbolide (100 μM final concentration, Cayman Chemical Co., 19230) or equivalent volume of DMSO. Samples were incubated at room temperature for 30 min. One microgram of pure p21 was added to each sample, and samples were incubated at room temperature 30 min with agitation. Ten microliters of Flag agarose beads were added to each sample, and samples were agitated at room temperature for 30 min. Washes (3 times, 1 mL TBS) were performed before proteins were eluted using 50 μL of TBS supplemented with 250 ng/μL 3 × FLAG peptide (ApexBio A6001). Supernatant (30 μL) were collected and after the addition of Laemmli’s reducing agent (10 μL), samples were boiled at 95 °C for 5 min and allowed to cool. Samples were analyzed by Western blotting as described above.

### *In situ* Nimbolide-Alkyne Probe Labeling and Biotin-Azide Pulldown

Experiments were performed following an adaption of a previously described protocol^81^. 231MFP cells were treated with either vehicle (DMSO) or 50 μM nimbolide alkyne probe for 90 min. Cells were harvested in PBS and lysed by sonication. For preparation of Western blotting sample, 195 μL of lysate was aliquoted per sample to which 25 μL of 10% SDS, 5 μL of 5 mM biotin picolylazide (900912 Sigma-Aldrich) and 25 μL of click reaction mix (3 parts TBTA 5 mM TBTA in butanol:DMSO (4:1, v/v), 1 part 50 mM Cu(II)SO4 solution, and 1 part 50 mM TCEP). Samples were incubated for 2 h at 37 °C with gentle agitation after which 1.2 mL ice cold acetone were added. After overnight precipitation at −20 °C, samples were spun in a prechilled centrifuge at 1250 g for 10 min allowing for aspiration of excess acetone and subsequent reconstitution of protein pellet in 200 μL PBS containing 1% SDS by probe sonication. At this stage, total protein concentration was determined by BCA assay and samples were normalized to a total volume of 230 μL, with 30 μL reserved as input. 20 μL of prewashed 50% streptavidin agarose bead slurry was added to each sample and samples were incubated overnight at rt with gentle agitation. Supernatant was aspirated from each sample after spinning at 90 g for 2 min at rt. Beads were transferred to spin columns and washed 3X with PBS. To elute, beads were boiled 5 min in 50 μL LDS sample buffer. Eluents were collected by centrifugation and analyzed by immunoblotting.

The resulting samples were analyzed as described below in the TMT-based quantitative proteomic profiling section.

### TMT-Based Quantitative Proteomic Profiling

#### Cell Lysis, Proteolysis and Isobaric Labeling

Treated cell-pellets were lysed and digested using the commercially available Pierce™ Mass Spec Sample Prep Kit for Cultured Cells (Thermo Fisher Scientific, P/N 84840) following manufacturer’s instructions. Briefly, 100 µg protein from each sample was reduced, alkylated, and digested overnight using a combination of Endoproteinase Lys-C and trypsin proteases. Individual samples were then labeled with isobaric tags using commercially available Tandem Mass Tag™ 6-plex (TMTsixplex™) (Thermo Fisher Scientific, P/N 90061) or TMT11plex (TMT11plex™) isobaric labeling reagent (Thermo Fisher Scientific, P/N 23275) kits, in accordance with manufacturer’s protocols.

#### High pH Reversed Phase Separation

Tandem mass tag labeled (TMT) samples were then consolidated, and separated using high-pH reversed phase chromatography (RP-10) with fraction collection as previously described^81^. Fractions were speed-vac dried, then reconstituted to produce 24 fractions for subsequent on-line nanoLC-MS/MS analysis.

#### Protein Identification and Quantitation by nanoLC-MS/MS

Reconstituted RP-10 fractions were analyzed on a Thermo Orbitrap Fusion Lumos Mass Spectrometer (Xcalibur 4.1, Tune Application 3.0.2041) coupled to an EasyLC 1200 HPLC system (Thermo Fisher Scientific). The EasyLC 1200 was equipped with a 20 μL loop, set-up for 96 well plates. A Kasil-fritted trapping column (75 µm ID) packed with ReproSil-Pur 120 C18-AQ, 5 µm material (15mm bed length) was utilized together with a 160mm length, 75 μm inner diameter spraying capillary pulled to a tip diameter of approximately 8-10 µm using a P-2000 capillary puller (Sutter Instruments, Novato, CA). The 160mm separation column was packed with ReproSil-Pur 120 C18-AQ, 3 µm material (Dr. Maisch GmbH, Ammerbuch-Entringen, Germany). Mobile phase consisted of A= 0.1% formic acid/2% acetonitrile (v/v), and Mobile phase B= 0.1% formic acid/98% acetonitrile (v/v). Samples (18 µL) were injected on to trapping column using Mobile Phase A at a flow rate of 2.5 μL/min. Peptides were then eluted using an 80 minute gradient (2% Mobile Phase B for 5 min, 2%-40% B from 5-65 min, followed by 70% B from 65-70 minutes, then returning to 2% B from 70-80 min) at a flowrate of 300 nL/min on the capillary separation column with direct spraying into the mass spectrometer. Data was acquired on Orbitrap Fusion Lumos Mass Spectrometer in data-dependent mode using synchronous precursor scanning MS^3^ mode (SPS-MS3), with MS^2^ triggered for the 12 most intense precursor ions within a mass-to-charge ratio *(m/z)* range of 300-1500 found in the full MS survey scan event. MS scans were acquired at 60,000 mass resolution (*R*) at *m/z* 400, using a target value of 4 × 10^5^ ions, and a maximum fill time of 50 ms. MS^2^ scans were acquired as CID ion trap (IT) rapid type scans using a target value of 1 × 10^4^ ions, maximum fill time of 50 ms, and an isolation window of 2 Da. Data-dependent MS^3^ spectra were acquired as Orbitrap (OT) scans, using Top 10 MS^2^ daughter selection, automatic gain control (AGC) target of 5 × 10^4^ ions, with scan range of *m/z* 100-500. The MS^3^ maximum injection time was 86 ms, with HCD collision energy set to 65%. MS^3^ mass resolution (*R*) was set to 15,000 at *m/z* 400 for TMT6plex experiments, and 50,000 at *m/z* 400 for TMT11-plex experiments. Dynamic exclusion was set to exclude selected precursors for 60 s with a repeat count of 1. Nanospray voltage was set to 2.2 kV, with heated capillary temperature set to 300 °C, and an S-lens RF level equal to 30%. No sheath or auxiliary gas flow is applied.

#### Data Processing and Analysis

Acquired MS data was processed using Proteome Discoverer v. 2.2.0.388 software (Thermo) utilizing Mascot v 2.5.1 search engine (Matrix Science, London, UK) together with Percolator validation node for peptide-spectral match filtering^82^. Data was searched against Uniprot protein database (canonical human and mouse sequences, EBI, Cambridge, UK) supplemented with sequences of common contaminants. Peptide search tolearances were set to 10 ppm for precursors, and 0.8 Da for fragments. Trypsin cleavage specificity (cleavage at K, R except if followed by P) allowed for up to 2 missed cleavages. Carbamidomethylation of cysteine was set as a fixed modification, methionine oxidation, and TMT-modification of N-termini and lysine residues were set as variable modifications. Data validation of peptide and protein identifications was done at the level of the complete dataset consisting of combined Mascot search results for all individual samples per experiment via the Percolator validation node in Proteome Discoverer. Reporter ion ratio calculations were performed using summed abundances with most confident centroid selected from 20 ppm window. Only peptide-to-spectrum matches that are unique assignments to a given identified protein within the total dataset are considered for protein quantitation. High confidence protein identifications were reported using a Percolator estimated <1% false discovery rate (FDR) cut-off. Differential abundance significance was estimated using a background-based ANOVA with Benjamini-Hochberg correction to determine adjusted p-values.

### Tumor Xenograft Studies

Human tumor xenografts were established by transplanting cancer cells ectopically into the flank of female C.B17 severe combined immunodeficiency (SCID) mice (Taconic Farms) as previously described^83, 84^. In brief, cells were washed with PBS, trypsinized, and harvested in serum-containing medium. Harvested cells were washed in serum-free media and resuspended in serum-free media at a concentration of 2.0 × 10^4^ cells/μl, and 100 μl was injected subcutaneously into the flank of each mouse. Tumors were measured with calipers. Animal experiments were conducted in accordance with the guidelines of the Institutional Animal Care and Use Committees of the University of California, Berkeley.

## Data Availability Statement

The datasets generated during and/or analyzed during the current study are available from the corresponding author on reasonable request.

## Acknowledgement

We thank the members of the Nomura Research Group, the Maimone lab, and Novartis Institutes for BioMedical Research for critical reading of the manuscript. We acknowledge Malte Moeller and Angus Olding for assistance in nimbolide isolation studies. This work was supported by Novartis Institutes for BioMedical Research and the Novartis-Berkeley Center for Proteomics and Chemistry Technologies (NB-CPACT) for all listed authors. This work was also supported by grants from the National Institutes of Health (R01CA172667 for DKN, JNS, CCW, LO; F31CA225173 for CCW; F31CA239327 for JNS).

