## Supporting Information for "Harnessing the Anti-Cancer Natural Product Nimbolide for Targeted Protein Degradation"

[7] Current address: Vertex Pharmaceuticals, Boston, MA 02210

### Supporting Methods

#### Synthesis and characterization of the nimbolide-alkyne probe (SI-2) and degraders XH1 and XH2

#### General Procedures

Unless otherwise stated, all reactions were performed in oven-dried or flame-dried Fisherbrand® borosilicate glass tubes (Fisher Scientific, 1495925A, 13 × 100 mm) with a black phenolic screw cap (13-425) under an atmosphere of dry nitrogen. Dry *N*,*N*-dimethylformamide (DMF), toluene, and acetonitrile were obtained by passing these previously degassed solvents through activated alumina columns. Nimbolide was purchased from Sigma-Aldrich or Cayman Chemical, and used directly without further purification. Propargylamine and *N*-methylpropargylamine were purchased from Fisher Scientific and used directly without further purification. Reactions were monitored by thin layer chromatography (TLC) on TLC silica gel 60 $F_{254}$ glass plates (EMD Millipore) and visualized by UV irradiation and staining with *p*-anisaldehyde, phosphomolybdic acid, or potassium permanganate. Volatile solvents were removed under reduced pressure using a rotary evaporator. Flash column chromatography was performed using Silicycle F60 silica gel (60Å, 230-400 mesh, 40-63 µm). Ethyl acetate and hexanes were purchased from Fisher Chemical and used for chromatography without further purification. Proton nuclear magnetic resonance ($^1$H NMR) and carbon nuclear magnetic resonance ($^{13}$C NMR) spectra were recorded on Bruker AV-600 and AV-700 spectrometers operating at 600 and 700 MHz for $^1$H, and 150 and 175 MHz for $^{13}$C. Chemical shifts are reported in parts per million (ppm) with respect to the residual solvent signal $CDCl_3$ ($^1$H NMR: δ = 7.26; $^{13}$C NMR: δ = 77.16), $CD_2Cl_2$ ($^1$H NMR: δ = 5.32; $^{13}$C NMR: δ = 53.84). Peak multiplicities are reported as follows: *s* = singlet, *d* = doublet, *t* = triplet, *dd* = doublet of doublets, *tt* = triplet of triplets, *m* = multiplet, *br* = broad signal, *app* = apparent. IR spectra were recorded on a Nicolet 380 FT-IR spectrometer. High-resolution mass spectra (HRMS) were obtained by the QB3/chemistry mass spectrometry facility at the University of California, Berkeley. Optical rotations were measured on a Perkin-Elmer 241 polarimeter.

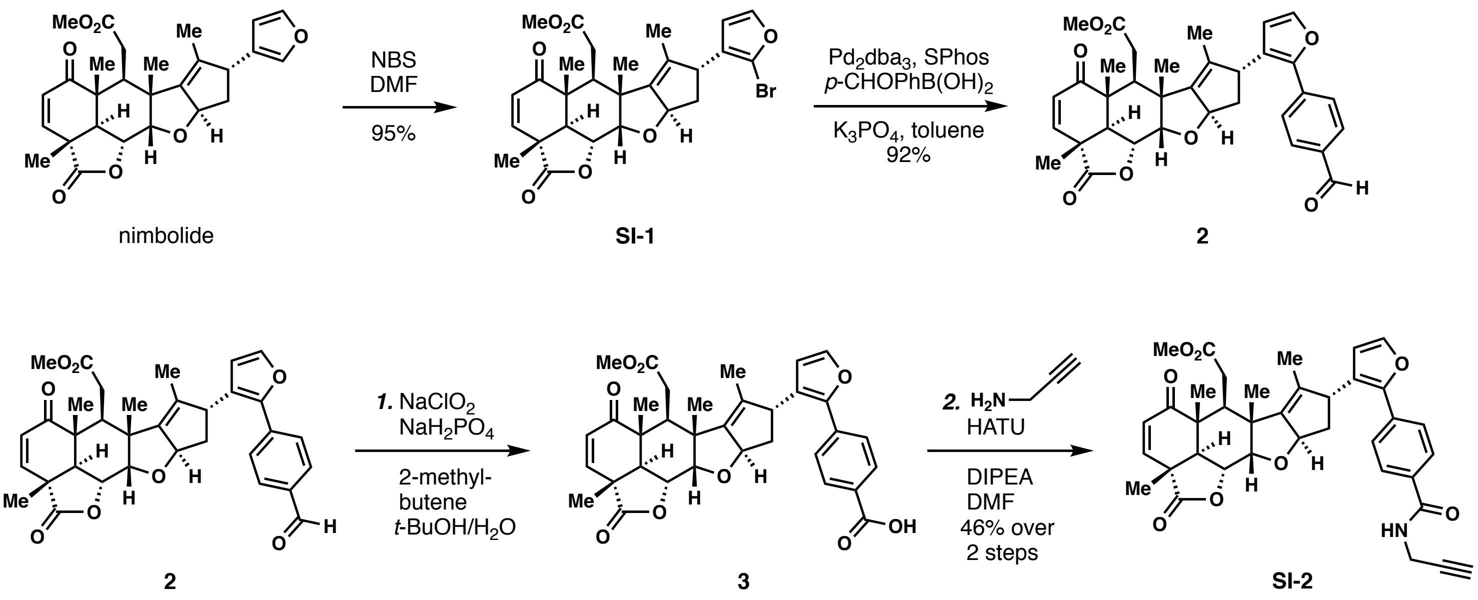


**Scheme S1.** Synthesis of the nimbolide alkyne probe (**SI-2**).

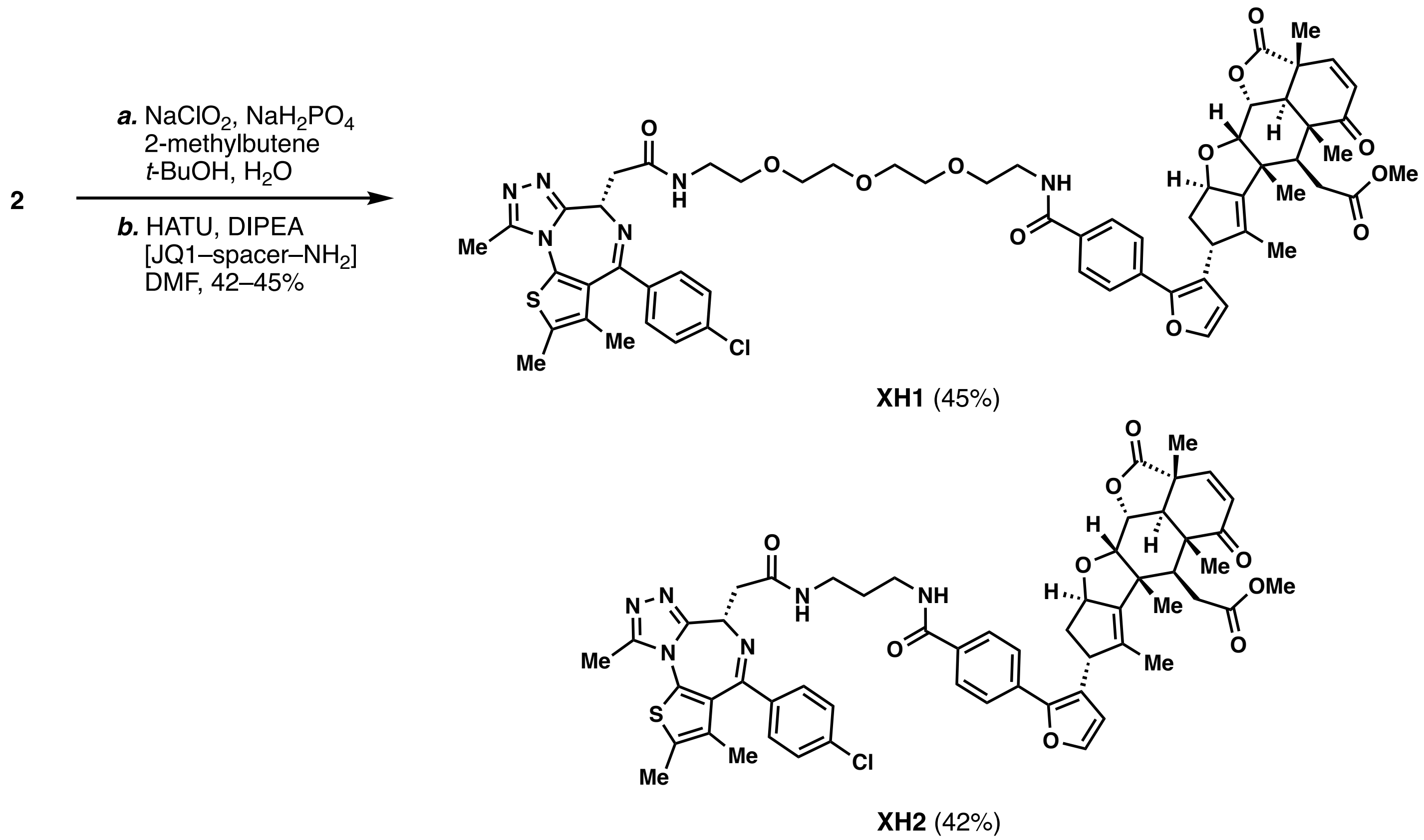


**Scheme S2.** Synthesis of nimbolide-derived bifunctional degraders **XH1 and XH2**.

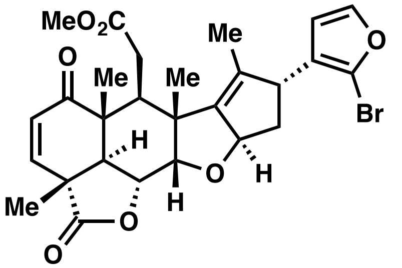

**Bromofuran SI-1**: Nimbolide (100 mg, 0.214 mmol) was divided evenly into four reaction tubes (Fisher Scientific, 13 × 100 mm), each charged with a stir bar. The tubes were evacuated and back-filled with nitrogen, dry DMF (0.25 mL each) was added, and the resulting solutions were cooled to 0 °C in an ice bath. Recrystallized *N*-bromosuccinimide (40.0 mg, 0.225 mmol) was dissolved in dry DMF (4 mL), and the solution was slowly added to each reaction tube (1 mL each). The reaction mixture was stirred at 0 °C for 1 hour and then quenched by the addition of saturated *aq*. $Na_2S_2O_3$ (5 mL each). The resulting mixtures were combined and extracted with EtOAc (3 X 20 mL). The combined organic layer was washed with $H_2O$ (50 mL) and brine (50 mL), dried over $MgSO_4$, and concentrated *in vacuo*. The crude mixture was purified by column chromatography (EtOAc:hexane = 1:3 to 1:1), affording **SI-1** (111 mg, 95%) as a white foam: $[\alpha]_D^{20}$ = +190.3° (c 0.010 g/ml, $CHCl_3$); $^1H$ NMR (700 MHz, $CDCl_3$) δ 7.34 (d, *J* = 2.1 Hz, 1H), 7.28 (d, *J* = 9.7 Hz, 1H), 6.29 (d, *J* = 2.1 Hz, 1H), 5.92 (d, *J* = 9.7 Hz, 1H), 5.56 (app. tt, *J* = 7.4, 1.8 Hz, 1H), 4.62 (dd, *J* = 12.5, 3.7 Hz, 1H), 4.27 (d, *J* = 3.7 Hz, 1H), 3.67 (brd, *J* = 7.0 Hz, 1H), 3.56 (s, 3H), 3.24 (dd, *J* = 16.3, 5.5 Hz, 1H), 3.18 (d, *J* = 12.5 Hz, 1H), 2.75 (dd, *J* = 5.5, 5.5 Hz, 1H), 2.36 (dd, *J* = 16.3, 5.5 Hz, 1H), 2.18 – 2.13 (m, 2H), 1.66 (d, *J* = 1.4 Hz, 3H), 1.47 (s, 3H), 1.36 (s, 3H), 1.22 (s, 3H); $^{13}C$ NMR (175 MHz, $CDCl_3$) δ 200.9, 175.0, 173.2, 149.8, 145.4, 144.2, 136.1, 131.2, 125.2, 120.3, 112.1, 88.5, 83.1, 73.5, 52.0, 50.5, 50.0, 47.9, 45.4, 43.8, 41.2, 40.4, 32.2, 18.7, 17.3, 15.3, 13.0; IR (thin film, $cm^{-1}$) 2974, 2929, 2875, 1783, 1734, 1678, 1594, 1438, 1394, 1373; HRMS (ESI) *calcd.* for $[C_{27}H_{29}O_7BrNa]^+$ $(M+Na)^+$: m/z 567.0989, found 567.0990.

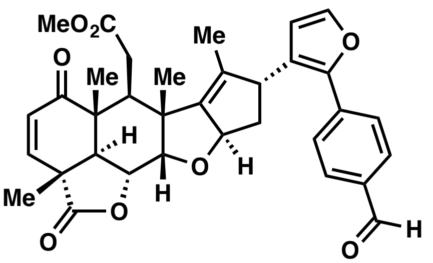

**Aldehyde 2**: A reaction tube (Fisher Scientific, 13 × 100 mm) was charged with a stir bar, **SI-1** (12 mg, 0.022 mmol, 1 equiv), $Pd_2(dba)_3$ (10 mg, 0.011 mmol, 0.5 equiv), SPhos (10 mg, 0.024 mmol, 1 equiv), anhydrous $K_3PO_4$ (35 mg, 0.17 mmol, 7.5 equiv) and 4-formylphenylboronic acid (16 mg, 0.11 mmol, 5 equiv). The tube was evacuated and back-filled with nitrogen and dry toluene (0.5 mL) added. The resulting mixture was heated at 60 ºC for 48 h, cooled to room temperature, and passed through a plug of Celite®. The filtrate was concentrated *in vacuo* and purified by column chromatography (EtOAc:hexane = 1:3 to 1:1) to afford aldehyde **2** (11.5 mg, 92%) as a light yellow oil: $[\alpha]_D^{20}$ = +4.8° (c 0.005 g/mL, $CHCl_3$); $^1$H NMR (600 MHz, $CDCl_3$) δ 10.01 (s, 1H), 7.92 (d, *J* = 8.3 Hz, 2H), 7.71 (d, *J* = 8.3 Hz, 2H), 7.42 (d, *J* = 1.9 Hz, 1H), 7.29 (d, *J* = 9.7 Hz, 1H), 6.39 (d, *J* = 1.9 Hz, 1H), 5.93 (d, *J* = 9.7 Hz, 1H), 5.60 (app. tt, *J* = 7.3, 1.7 Hz, 1H), 4.64 (dd, *J* = 12.5, 3.6 Hz, 1H), 4.32 (d, *J* = 3.6 Hz, 1H), 4.17 – 4.13 (m, 1H), 3.69 (s, 3H), 3.23 (dd, *J* = 16.4, 5.5 Hz, 1H), 3.20 (d, *J* = 12.5 Hz, 1H), 2.78 (dd, *J* = 5.5, 5.5 Hz, 1H), 2.41 (dd, *J* = 16.4, 5.5 Hz, 1H), 2.31 – 2.27 (m, 2H), 1.69 (d, *J* = 1.7 Hz, 3H), 1.49 (s, 3H), 1.38 (s, 3H), 1.25 (s, 3H); $^{13}$C NMR (150 MHz, $CDCl_3$) δ 200.8, 191.7, 175.0, 173.2, 149.8, 147.9, 146.3, 143.1, 137.0, 136.1, 134.8, 131.1, 130.3, 126.1, 126.0, 112.6, 88.4, 83.3, 73.4, 52.1, 50.8, 50.0, 47.9, 45.5, 43.8, 41.3, 32.5, 18.8, 17.5, 15.3, 13.4; IR (thin film, $cm^{-1}$) 2977, 2935, 2873, 1782, 1733, 1702, 1677, 1608, 1438, 1393, 1372; HRMS (ESI) *calcd.* for $[C_{34}H_{34}O_8Na]^+$ $(M+Na)^+$: m/z 593.2146, found 593.2154.
[Note: Aldehyde **2** is not stable under preparative TLC conditions]

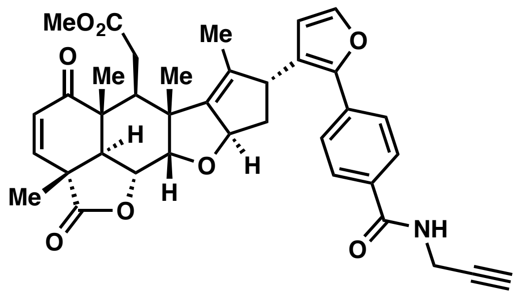

**Nimbolide Alkyne Probe SI-2:** *i.* A reaction tube (Fisher Scientific, 13 × 100 mm) was charged with a stir bar, aldehyde **2** (7.0 mg, 0.012 mmol, 1 equiv), and a mixture of *t*-BuOH and 2-methyl-2-butene (0.6 mL, 3:5 v/v). A solution of $NaClO_2$ (3.3 mg, 3 equiv) and $NaH_2PO_4$ (13.2 mg, 9 equiv) in $H_2O$ (0.2 mL) was added in one portion and the resulting mixture was stirred at room temperature for 6 hours. After the reaction was complete as judged by TLC (EtOAc:hexane = 2:1), the mixture was diluted with EtOAc (10 mL) and saturated *aq.* $NH_4Cl$ (10 mL), and the aqueous phase was extracted by EtOAc (10 mL × 2). The combined organic layer was washed with brine (20 mL), dried over $MgSO_4$, and concentrated *in vacuo*. The resulting crude material was used directly without further purification.
*ii.* A reaction tube (Fisher Scientific, 13 × 100 mm) charged with a stir bar, crude **3** (0.012 mmol assumed), and HATU (13.7 mg, 0.036 mmol, 3 equiv) was added a solution of DIPEA (6.4 μL, 0.036 mmol, 3 equiv) in DMF (0.1 mL). The resulting mixture was cooled to 0 ºC and stirred for 10 minutes. A solution of propargylamine (1.6 μL, 0.024 mmol, 2 equiv) in DMF (0.2 mL) was then added and the reaction mixture was further stirred at 0–4 ºC for 12 hours. After the reaction was complete, as judged by TLC (EtOAc:hexane = 2:1), the mixture was diluted with EtOAc (10 mL) and saturated *aq.* $NH_4Cl$ (10 mL), and the aqueous phase was extracted by EtOAc (10 mL × 2). The combined organic layer was washed with $H_2O$ (20 mL), brine (20 mL), dried over $MgSO_4$, and concentrated *in vacuo*. The resulting crude was purified by preparative TLC (EtOAc:hexane = 2:1), affording **SI-3** (3.5 mg, 46% over 2 steps) as a white solid: $[\alpha]_D^{20}$ = +21.5° (c 0.002 g/mL, $CHCl_3$); $^1$H NMR (600 MHz, $CDCl_3$) δ 7.83 (d, *J* = 8.2 Hz, 2H), 7.61 (d, *J* = 8.2 Hz, 2H), 7.39 (d, *J* = 1.8 Hz, 1H), 7.29 (d, *J* = 9.7 Hz, 1H), 6.36 (d, *J* = 1.8 Hz, 1H), 6.27 (t, *J* = 5.2 Hz, 1H), 5.93 (d, *J* = 9.7 Hz, 1H), 5.60 (app. t, *J* = 7.6 Hz, 1H), 4.64 (dd, *J* = 12.6, 3.7 Hz, 1H), 4.31 (d, *J* = 3.7 Hz, 1H), 4.28 (dd, *J* = 5.2, 2.6 Hz, 2H), 4.11 (brd, *J* = 7.6 Hz, 1H), 3.68 (s, 3H), 3.25 – 3.18 (m, 2H), 2.78 (dd, *J* = 5.5, 5.5 Hz, 1H), 2.40 (dd, *J* = 16.4, 5.5 Hz, 1H), 2.32 – 2.24 (m, 3H), 1.67 (brs, 3H), 1.49 (s, 3H), 1.38 (s, 3H), 1.24 (s, 3H); $^{13}$C NMR (150 MHz, $CDCl_3$) δ 200.8, 175.0, 173.2, 166.6, 149.8, 148.2, 146.1, 142.5, 136.3, 134.7, 132.1, 131.2, 127.6, 126.1, 124.8, 112.3, 88.5, 83.2, 79.6, 73.5, 72.2, 52.1, 50.8, 50.0, 48.0, 45.5, 43.8, 41.3, 41.3, 32.5, 30.0, 18.8, 17.5, 15.3, 13.4; IR (thin film, $cm^{-1}$) 3337, 2954, 2921, 2852, 1781, 1734, 1663, 1609; HRMS (ESI) *calcd.* for $[C_{37}H_{37}NO_8Na]^+$ $(M+Na)^+$: m/z 646.2417, found 646.2417.

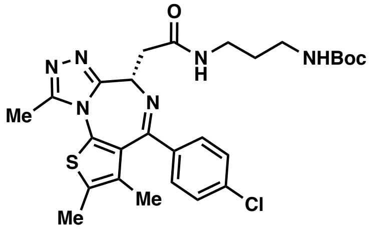

This compound was prepared according to conditions reported by Waring and coworkers (*J. Med. Chem.*, **2016**, *59*, 7801).

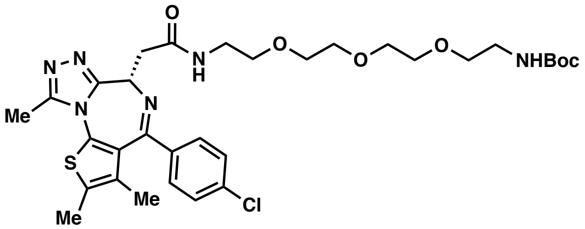

This compound was prepared according to the conditions reported by Bradner and co-workers (PCT Int. Appl. 2017, WO 2017091673 A2).

**General procedure for the synthesis of bifunctional degraders XH1 and XH2:**

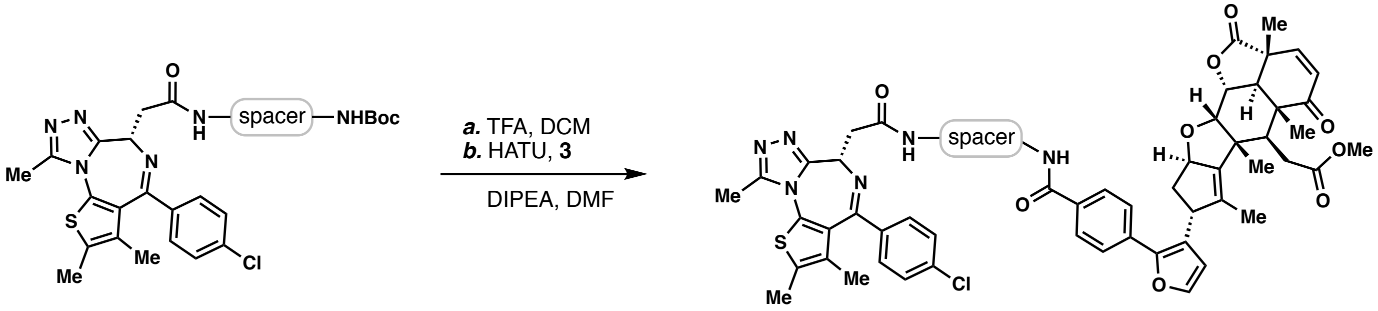


*i.* A reaction tube (Fisher Scientific, 13 × 100 mm) was charged with a stir bar, the Boc-protected amine (0.04 mmol), triethylsilane (12 μL, 0.075 mmol), and DCM (0.4 mL). The resulting mixture was cooled to 0 ºC, followed by the dropwise addition of TFA (0.12 mL). The reaction mixture was allowed to warm to room temperature and further stirred for 2 hours. After the reaction was complete as judged by TLC (MeOH:DCM = 1:10), the mixture was diluted with toluene (2 mL), and concentrated *in vacuo*. The resulting crude was dried under high vacuum for 30 minutes, and directly used in the next step without further purification. Acid **3** was prepared from aldehyde **2** according to the aforementioned procedure, and used without further purification.

*ii.* A reaction tube (Fisher Scientific, 13 × 100 mm) was charged with a stir bar, the crude amine (0.04 mmol assumed), unpurified **3** (0.02 mmol assumed), and DMF (0.6 mL). The resulting mixture was cooled to 0 ºC in an ice bath, followed by the addition of HATU (23.0 mg, 0.06 mmol) and DIPEA (11 μL, 0.06 mmol). The reaction mixture was stirred at 4 ºC for 16 hours. After the reaction was complete as judged by TLC (MeOH:DCM = 1:10), the mixture was diluted with EtOAc (10 mL) and saturated *aq.* $NH_4Cl$ (10 mL), and the aqueous phase extracted with EtOAc (2 x 10 mL). The combined organic layer was washed with $H_2O$ (20 mL), brine (20 mL), dried over $MgSO_4$, and concentrated *in vacuo*. The resulting crude material was purified by preparative TLC (MeOH:DCM = 1:15, developed twice), affording the bifunctional degraders **XH1** or **XH2**.

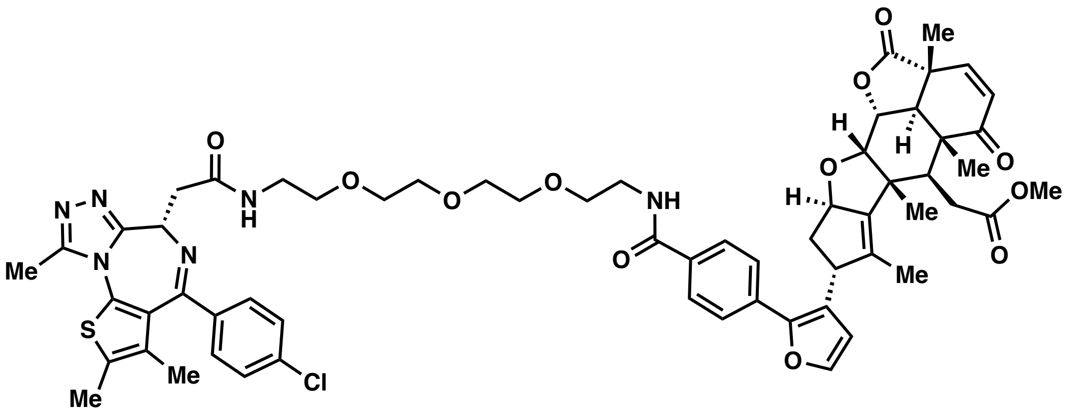

**XH1** (45% yield from **SI-2**), a white foam: $[\alpha]_D^{20}$ = +48.7° (c 0.0052 g/mL, $CHCl_3$); $^1H$ NMR (600 MHz, $CDCl_3$) δ 7.87 (d, *J* = 8.5 Hz, 2H), 7.53 (d, *J* = 8.5 Hz, 2H), 7.40 (d, *J* = 8.2 Hz, 3H), 7.36 (d, *J* = 1.9 Hz, 1H), 7.34 – 7.29 (m, 2H), 7.28 (d, *J* = 9.7 Hz, 1H), 6.33 (d, *J* = 1.9 Hz, 1H), 5.93 (d, *J* = 9.7 Hz, 1H), 5.61 – 5.55 (m, 1H), 4.73 (app. t, *J* = 7.1 Hz, 1H), 4.64 (dd, *J* = 12.5, 3.7 Hz, 1H), 4.30 (d, *J* = 3.6 Hz, 1H), 4.12 – 4.08 (m, 1H), 3.75 – 3.64 (m, 15H), 3.64 – 3.53 (m, 3H), 3.51 – 3.40 (m, 3H), 3.25 – 3.16 (m, 2H), 2.77 (app. t, *J* = 5.5 Hz, 1H), 2.70 (s, 3H), 2.42 – 2.36 (m, 4H), 2.26 – 2.20 (m, 2H), 1.68 – 1.65 (m, 3H), 1.65 (d, *J* = 1.8 Hz, 3H), 1.48 (s, 3H), 1.37 (s, 3H), 1.24 (s, 3H); $^{13}C$ NMR (150 MHz, $CDCl_3$) δ 200.6, 174.8, 173.0, 170.1, 167.0, 164.4, 155.3, 149.9, 149.6, 148.3, 146.1, 145.6, 142.1, 137.2, 136.3, 135.8, 133.8, 133.0, 131.6, 131.2, 131.0, 130.6, 130.0, 130.0, 128.7, 128.7, 127.6, 127.6, 125.7, 125.7, 124.1, 112.0, 88.3, 83.0, 73.3, 70.5, 70.5, 70.3, 70.2, 69.7, 69.6, 54.1, 51.9, 50.5, 49.7, 47.8, 45.3, 43.6, 41.1, 41.1, 39.7, 39.4, 38.5, 32.3, 18.6, 17.3, 15.1, 14.4, 13.1, 13.1, 11.6; IR (thin film, $cm^{-1}$) 3339, 2923, 2866, 1781, 1734, 1659, 1540, 1487, 1437, 1419, 1300; HRMS (ESI) *calcd.* for $[C_{61}H_{67}N_6O_{12}Cl_1S_1Na]^+$ $(M+Na)^+$: m/z 1165.4118, found 1165.4142.

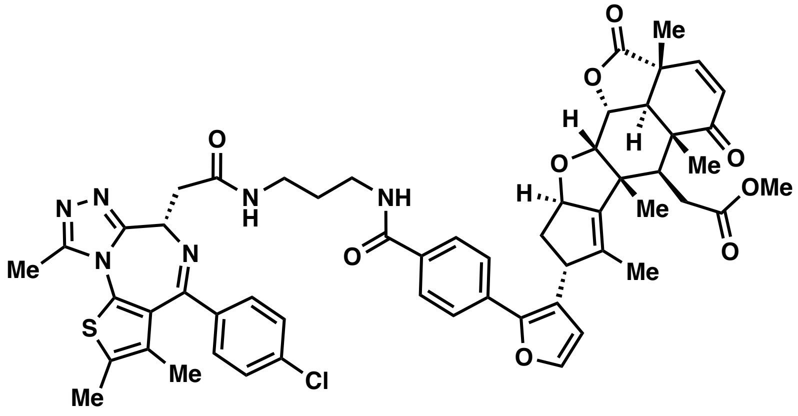

**XH2** (42% yield from **SI-2**), a white foam: $[\alpha]_D^{20}$ = +18.6° (c 0.004 g/mL, $CHCl_3$); $^1H$ NMR (600 MHz, $CD_2Cl_2$) δ 7.90 (d, *J* = 8.3 Hz, 2H), 7.65 – 7.57 (m, 3H), 7.44 – 7.39 (m, 3H), 7.33 (d, *J* = 8.5 Hz, 2H), 7.25 (d, *J* = 9.7 Hz, 1H), 7.07 (t, *J* = 6.5 Hz, 1H), 6.36 (d, *J* = 1.9 Hz, 1H), 5.89 (d, *J* = 9.7 Hz, 1H), 5.52 (app. t, *J* = 7.6 Hz, 1H), 4.66 – 4.59 (m, 2H), 4.26 (d, *J* = 3.6 Hz, 1H), 4.12 (brd, *J* = 7.6 Hz, 1H), 3.63 (s, 3H), 3.52 – 3.45 (m, 2H), 3.44 – 3.34 (m, 4H), 3.20 (dd, *J* = 16.4, 5.5 Hz, 1H), 3.15 (d, *J* = 12.5 Hz, 1H), 2.74 (app. t, *J* = 5.5 Hz, 1H), 2.63 (s, 3H), 2.45 – 2.37 (m, 4H), 2.25 – 2.21 (m, 2H), 1.75 – 1.70 (m, 2H), 1.65 (s, 6H), 1.46 (s, 3H), 1.36 (s, 3H), 1.22 (s, 3H); $^{13}C$ NMR (150 MHz, $CD_2Cl_2$) δ 201.2, 175.6, 173.5, 171.9, 166.7, 164.5, 156.1, 150.5, 149.9, 148.8, 146.3, 142.5, 137.1, 137.0, 136.6, 134.2, 133.7, 132.7, 131.6, 131.3, 131.2, 130.8, 130.3, 130.3, 129.0, 129.0, 127.8, 127.8, 126.2, 126.2, 124.8, 112.6, 88.6, 83.4, 73.9, 54.9, 52.1, 51.0, 50.2, 48.2, 45.8, 44.1, 41.6, 41.5, 39.7, 36.5, 36.2, 32.7, 29.9, 18.8, 17.5, 15.4, 14.6, 13.4, 13.3, 12.0; IR (thin film, $cm^{-1}$) 3308, 3023, 2930, 2867, 1782, 1734, 1654, 1540, 1488, 1437, 1301; HRMS (ESI) *calcd.* for $[C_{56}H_{57}ClN_6O_9S]^+$ $(M+H)^+$: m/z 1025.3669, found 1025.3669.

**Synthesis and characterization of JNS27**

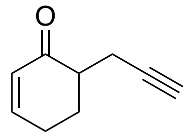

**JNS27.** i. To a stirring solution of lithium diisopropylamide (6.0 mL, 1.2 equiv) at -78 °C under nitrogen atmosphere, was added a solution of cyclohexenone (0.48mL, 1 equiv) in THF (5.0 mL) drop wise. After 45 minutes, a solution of 3-bromo-1-(trimethylsilyl)-propyne (0.85 mL, 2.4 equiv) in THF (5.0 mL) was added gradually, before allowing the reaction to come to room temperature overnight. The reaction was quenched with saturated $NH_4Cl$ (20mL) and extracted with EtOAc (20 mL × 3). The combined organic layers were washed with brine, dried with $Na_2SO_4$, and concentrated *in vacuo*. Flash chromatography of the crude residue on silica gel (10-20% EtOAc in hexanes) gave the TMS-protected alkyne in 25% yield (261 mg). ii. To a solution of this material (261 mg, 1 equiv) in THF (5.0 mL) under $N_2$ was added a solution of TBAF (0.44 mL, 1 equiv). After 4 hours of stirring at room temperature, the reaction was quenched by the addition of saturated $NH_4Cl$ (15 mL). The mixture was extracted with EtOAc (3 x15 mL). The combined organic layers were washed with brine, dried with $Na_2SO_4$, and concentrated *in vacuo*. Flash chromatography of the residue on silica gel (0-20% EtOAc in hexanes) gave JNS-27 as a pale-yellow waxy solid in 81% yield (168 mg, 21% over 2 steps). $^1$H NMR (400 MHz, $CDCl_3$) δ 6.99 (m, 1H), 6.03 (dt, *J* = 10.02, 1.97 Hz, 1H), 2.78 (m, 1H), 2.48 (m, 3H), 2.33 (m, 2H), 1.98 (t, *J* = 2.68, 1H), 1.88 (m, 1H); $^{13}$C NMR (400 MHz, $CDCl_3$) δ 150.30, 129.34, 45.74, 27.68, 25.78, 20.27; HRMS (ESI) *calcd.* for $[C_9H_{10}ONa]^+$ $(M+Na)^+$: m/z 134.0732, found 134.0728.

**Synthesis and characterization of cysteine-reactive covalent ligands previously not reported**
**General synthetic methods**

Chemicals and reagents were purchased from major commercial suppliers and used without further purification. Reactions were performed under a nitrogen atmosphere unless otherwise noted. Silica gel flash column chromatography was performed using EMD or Sigma Aldrich silica gel 60 (230-400 mesh). Proton and carbon nuclear magnetic resonance ($^1$H NMR and $^{13}$C NMR) data was acquired on a Bruker AVB 400, AVQ 400, or AV 600 spectrometer at the University of California, Berkeley. High resolution mass spectrum were obtained from the QB3 mass spectrometry facility at the University of California, Berkeley using positive or negative electrospray ionization (+ESI or -ESI). Yields are reported as a single run.

**General Procedure A**

The amine (1 eq.) was dissolved in DCM (5 mL/mmol) and cooled to 0$^\circ$C. To the solution was added acryloyl chloride (1.2 eq.) followed by triethylamine (1.2 eq.). The solution was warmed to room temperature and stirred overnight. The solution was then washed with brine and the crude product was purified by silica gel chromatography (and recrystallization if necessary) to afford the corresponding acrylamide.

**General Procedure B**

The amine (1 eq.) was dissolved in DCM (5 mL/mmol) and cooled to 0$^\circ$C. To the solution was added chloroacetyl chloride (1.2 eq.) followed by triethylamine (1.2 eq.). The solution was warmed to room temperature and stirred overnight. The solution was then washed with brine and the crude product was purified by silica gel chromatography (and recrystallization if necessary) to afford the corresponding chloroacetamide.

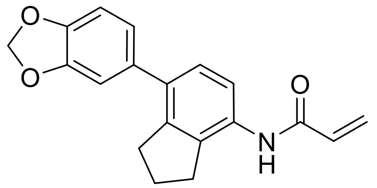

***N*-(7-(benzo[*d*][1,3]dioxol-5-yl)-2,3-dihydro-1*H*-inden-4-yl)acrylamide (TRH 1-78).**

To a solution of *N*-(7-bromo-2,3-dihydro-1H-inden-4yl)acrylamide (**TRH 1-65**, 55 mg, 0.2 mmol) in a mixture of dioxane and water (4:1 *v:v*, 2.1 mL) under nitrogen atmosphere was added sequentially 3,4-(methylenedioxy)phenylboronic acid (70 mg, 0.4 mmol), potassium carbonate (74 mg, 0.5 mmol), and tetrakis(triphenylphosphine)palladium(0) (24 mg, 10 mol%). The reaction mixture was heated to reflux and stirred overnight. The reaction was diluted with water (20 mL) and extracted with DCM (3x20 mL). The combined organics were dried with magnesium sulfate, filtered, and evaporated *in vacuo*. The resulting crude material was purified by silica gel chromatography (0% to 25% ethyl acetate in hexanes) to afford the product (7 mg, 11% yield) as a white solid.

**$^{1}$H NMR (600 MHz, $CDCl_3$)**: δ 7.93 (d, *J* = 7.0 Hz, 1H), 7.18 (d, *J* = 8.2 Hz, 1H), 7.10 (s, 1H), 6.90 (s, 1H), 6.86 (t, *J* = 8.1 Hz, 2H), 6.45 (d, *J* = 16.8 Hz, 1H), 6.30 (dd, *J* = 10.3, 16.8 Hz, 1H), 5.79 (d, *J* = 10.2 Hz, 1H), 3.00 (t, *J* = 7.3 Hz, 2H), 2.88 (t, *J* = 7.3 Hz, 2H), 2.10 (quint, 7.3 Hz, 2H).
**$^{13}$C NMR (150 MHz, $CDCl_3$)**: δ 147.7, 146.7, 142.84, 142.81, 135.2, 134.9, 132.8, 131.3, 127.9, 127.8, 122.1, 119.8, 109.2, 108.3, 101.2, 36.8, 33.5, 30.5, 25.4.
**HRMS (-ESI)**: Calculated: 306.1136 ($C_{19}H_{16}NO_3$). Observed: 306.1130.

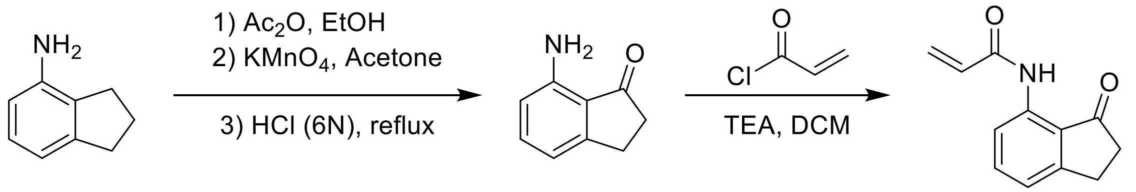


***N*-(3-oxo-2,3-dihydro-1*H*-inden-4-yl)acrylamide (TRH 1-129)**

i. To a solution of 4-aminoindan (1.0 g, 7.5 mmol) in ethanol (20 mL) at 0 °C was added acetic anhydride (1.4 mL, 15.0 mmol). The solution was then warmed to room temperature and stirred overnight at which point the solvent was evaporated *in vacuo*. The residue was then dissolved in acetone (50 mL) and 15% aqueous magnesium sulfate (1.2 g in 6.75 mL of water) followed by potassium permanganate (3.4 g, 17.0 mmol) were added. The resulting solution was stirred for 24 hours and then filtered through a pad of celite, eluting with chloroform and then water. The eluent was separated, and the aqueous layer was extracted several times with additional chloroform. The combined organics were dried over magnesium sulfate, filtered and evaporated *in vacuo*. The residue was then dissolved in 6N HCl (20 mL) and heated to 90 °C. After stirring for 5 hours, the solution was cooled, neutralized with small portions of potassium carbonate, and extracted with ethyl acetate. The combined organics were dried with magnesium sulfate, filtered, and evaporated *in vacuo* to give the crude **7-aminoindan-1-one** (610 mg, 55% over 3 steps) which was used without further purification.

ii. To a solution of the crude 7-aminoindan-1-one in dichloromethane (15 mL) was added acryloyl chloride (0.39 mL, 4.8 mmol) followed by triethylamine (0.67 mL, 4.8 mmol) at 0˚C under an atmosphere of $N_2$. The reaction mixture was warmed to room temperature and stirred overnight. The reaction mixture was then washed with 1M HCl solution twice, brine, and concentrated *in vacuo*. The crude material was purified by silica gel chromatography (10% to 20% ethyl acetate in hexanes) to yield the product (390 mg, 47% yield, 26% combined yield over 4 steps) as a white solid.

**$^{1}$H NMR (400MHz, $CDCl_3$)**: δ 10.64 (s, 1H), 8.45 (d, *J* = 8.2 Hz, 1H), 7.55 (t, *J* = 7.9 Hz, 1H), 7.12 (d, *J* = 7.6 Hz, 1H), 6.45 (dd, *J* = 1.0, 17.0 Hz, 1H), 6.33 (dd, *J* = 10.1, 17.0 Hz, 1H), 5.82 (dd, *J* = 1.0, 10.1 Hz, 1H), 3.11 (t, *J* = 11.5 Hz, 2H), 2.74-2.71 (m, 2H).
**$^{13}$C NMR (100MHz, $CDCl_3$)**: δ 209.3, 164.4, 155.9, 138.7, 137.0, 131.7, 128.0, 123.1, 120.8, 116.9, 36.5, 25.5.
**HRMS (+ESI)**: Calculated: 202.0863 ($C_{12}H_{12}NO_2$). Observed: 202.0860.

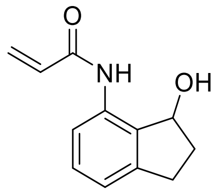

***N*-(3-hydroxy-2,3-dihydro-1*H*-inden-4-yl)acrylamide (TRH 1-133).**
To a solution of *N*-(3-oxo-2,3-dihydro-1*H*-inden-4-yl)acrylamide (**TRH 1-129**, 201 mg, 1.0 mmol) in anhydrous methanol (7 mL) under nitrogen atmosphere was added sodium borohydride (46.1 mg, 1.2 mmol). After 30 minutes of stirring, the reaction was quenched with saturated sodium bicarbonate solution and extracted three times with DCM. The combined organic layers were dried with magnesium sulfate, filtered, and concentrated *in vacuo*. The crude material was purified by silica gel chromatography (30 to 50% ethyl acetate in hexanes) affording the product (190 mg, 94% yield) as a white solid.
**$^{1}$H NMR (400 MHz, CDCl$_3$)**: δ 8.93 (s, 1H), 7.98 (d, *J* = 7.8 Hz, 1H), 7.19 (t, *J* = 7.9 Hz, 1H), 6.95 (d, *J* = 7.4 Hz, 1H), 6.29 (d, *J* = 16.8 Hz, 1H), 6.15 (dd, *J* = 10.2, 16.9 Hz, 1H), 5.66 (d, *J* = 10.2 Hz, 1H), 5.32 (q, *J* = 6.9 Hz, 1H), 3.60 (d, *J* = 6.7 Hz, 1H), 2.96 (ddd, *J* = 2.4, 9.0, 15.7 Hz), 2.73 (quint, *J* = 8.1 Hz, 1H), 2.56-2.48 (m, 1H), 1.96-1.86 (m, 1H).
**$^{13}$C NMR (100 MHz, CDCl$_3$)**: δ 164.1, 143,7 ,135.6, 132.8, 131.6, 129.5, 127.3, 121.0, 118.5, 76.2, 36.0, 29.8.
**HRMS (-ESI)**: Calculated: 202.0874 ($C_{12}H_{12}NO_2$). Observed: 202.0874.

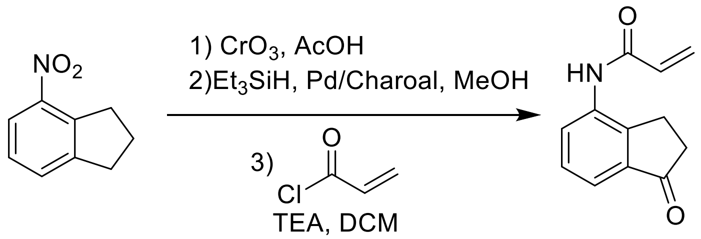


***N*-(1-oxo-2,3-dihydro-1*H*-inden-4-yl)acrylamide (TRH 1-134).**
*i.* To a solution of 4-nitroindan (5.38 g, 33 mmol) in acetic acid (250 mL) was slowly added chromium trioxide (8.95 g, 90 mmol). After stirring for 24 hours, the reaction was neutralized with 2M NaOH and extracted five times with ethyl acetate. The combined organics were washed with a saturated sodium bicarbonate solution and brine, dried over magnesium sulfate, filtered, and concentrated *in vacuo*. The crude material was purified by silica gel chromatography (10-20% ethyl acetate in hexanes) to give 1.26 g (ca. 7.1 mmol) of 4-nitroindanone as a white solid.
*ii.* This intermediate was then combined with palladium on activated charcoal (125 mg, 10 wt%) dissolved in anhydrous methanol (21 mL) under an atmosphere of a nitrogen. Triethylsilane (11.3 mL, 71 mmol) was slowly added by addition funnel over the course of 10 minutes to the reaction in a room temperature water bath. After an additional 20 minutes of stirring, the reaction mixture was filtered through a pad of celite and concentrated *in vacuo* to give 4-aminoindanone which was used without further purification.
*iii.* The aforementioned crude aminoindanone was dissolved in DCM (21 mL) under an atmosphere of nitrogen and cooled to 0 ˚C at which point acryloyl chloride (0.77 mL, 9.5 mmol) and triethylamine (1.19 mL, 8.5 mmol) were slowly added dropwise. The reaction mixture was warmed to room temperature, stirred overnight, washed twice with brine, dried over magnesium sulfate, filtered, and concentrated *in vacuo*. The crude material was purified by silica gel chromatography (30-50% ethyl acetate in hexanes) to give the title compound (989 mg, 15% yield over 3 steps) as a white solid.
**$^{1}$H NMR (400 MHz, CDCl$_3$)**: δ 8.20 (d, *J* = 5.8 Hz, 1H), 7.63 (s, 1H), 7.56 (d, *J* = 7.5 Hz, 1H), 7.39 (t, *J* = 7.7 Hz, 1H), 6.48 (d, *J* = 16.7 Hz, 1H), 6.37 (dd, *J* = 10.0 Hz, 16.8 Hz, 1H), 5.83 (d, *J* = 10.1 Hz, 1H), 3.04 (t, *J* = 5.6 Hz, 2H), 2.70 (t, *J* = 5.7 Hz, 2H).
**$^{13}$C NMR (100 MHz, CDCl$_3$)**: δ 206.3, 163.9, 146.0, 138.0, 135.4, 130.7, 128.8, 128.7, 127.6, 120.4, 36.1, 23.4.
**HRMS (-ESI)**: Calculated: 200.0717 ($C_{12}H_{10}NO_2$). Observed: 200.0715.

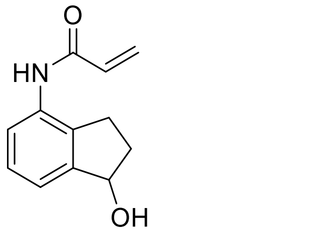


***N*-(1-hydroxy-2,3-dihydro-1*H*-inden-4-yl)acrylamide (TRH 1-135).**
To a solution of *N*-(1-oxo-2,3-dihydro-1*H*-inden-4-yl)acrylamide (**TRH 1-134**, 1.26 g, 6.25 mmol) in anhydrous methanol (50 mL) under nitrogen atmosphere was added sodium borohydride (292.7 mg, 7.7 mmol). After 30 minutes of stirring, the reaction was quenched with water and the methanol was removed *in vacuo*. The residue was saturated with NaCl and extracted five times with a 2:1 chloroform:methanol solution. The combined organic layers were dried over 3Å molecular sieves, filtered, and concentrated *in vacuo*. The crude material was purified by silica gel chromatography (40 to 70% ethyl acetate in hexanes) to give the product (1.05 g, 83% yield) as a white solid.
**$^{1}$H NMR (400 MHz, MeOD)**: δ 7.50 (dd, *J* = 2.3, 6.3 Hz, 1H), 7.25-7.20 (m, 2H), 6.51 (dd, *J* = 10.2, 17.0 Hz, 1H), 6.35 (dd, *J* = 1.7, 17.0 Hz, 1H), 5.77 (dd, *J* = 1.7, 10.2 Hz, 1H), 5.17 (t, *J* = 6.3 Hz, 1H), 2.97 (ddd, *J* = 4.5, 8.6, 16.2, 1H), 2.74 (quint, *J* = 7.8 Hz, 1H), 2.47-2.39 (m, 1H), 1.95-1.86 (m, 1H).
**$^{13}$C NMR (100 MHz, MeOD)**: δ 166.3, 148.0, 137.8, 134.8, 132.1, 128.3, 127.9, 124.0, 122.7, 76.9, 36.1, 28.6.
**HRMS (-ESI)**: Calculated: 202.0874 ($C_{12}H_{12}NO_2$). Observed: 202.0872.

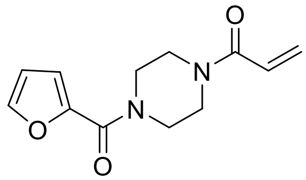


**1-(4-(furan-2-carbonyl)piperazin-1-yl)prop-2-en-1-one (TRH 1-145).**
To a solution 1-(2-furoyl)piperazine (362 mg, 2.0 mmol) in dichloromethane (10 mL) was added acryloyl chloride (0.20 mL, 2.4 mmol) followed by triethylamine (0.34 mL, 2.4 mmol) at 0˚C under an atmosphere of nitrogen. After stirring for 20 minutes, the reaction mixture was warmed to room temperature and was stirred an additional 24 hours. The reaction mixture was washed twice with brine, dried over magnesium sulfate, and concentrated *in vacuo*. The resulting crude material was purified by silica gel chromatography (70% to 100% ethyl acetate in hexanes) to yield the product (446 mg, 95%) as a yellow solid.
**$^{1}$H NMR (400 MHz, $CDCl_3$)**: δ 7.53 (m, 1H), 7.06 (dd, *J* = 0.7, 3.5 Hz, 1H), 6.61 (dd, *J* = 10.5, 16.8 Hz, 1H), 6.52 (dd, *J* = 1.8, 3.5 Hz, 1H), 6.33 (dd, *J* = 1.9, 16.8 Hz, 1H), 5.75 (dd, *J* = 1.9, 10.5 Hz, 1H), 3.84-3.67 (m, 8H).
**$^{13}$C NMR (100 MHz, $CDCl_3$)**: δ 165.5, 159.1, 147.5, 144.0, 128.5, 127.1, 117.0, 111.5, 45.6, 41.9.
**HRMS (+ESI)**: Calculated: 235.1077 ($C_{12}H_{15}N_2O_3$). Observed: 235.1075.

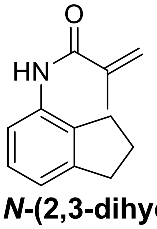


***N*-(2,3-dihydro-1*H*-inden-4-yl)methacrylamide (TRH 1-149).**
To a solution 4-aminoindan (0.24 mL, 2.0 mmol) in dichloromethane (10 mL) was added methacryloyl chloride (0.23 mL, 2.4 mmol) followed by triethylamine (0.34 mL, 2.4 mmol) at 0˚C under an atmosphere of nitrogen. After stirring for 20 minutes, the reaction mixture was warmed to room temperature and stirred for an additional 3.5 hours. The reaction mixture was then washed twice with brine, dried over magnesium sulfate, and concentrated *in vacuo*. The resulting crude material was purified by silica gel chromatography (35% to 40% ethyl acetate in hexanes) to yield the title compound (378 mg, 94%) as an off-white solid.
**$^{1}$H NMR (400 MHz, $CDCl_3$)**: δ 7.72 (d, *J* = 8.0 Hz, 1H), 7.55 (s, 1H), 7.12 (t, *J* = 7.7 Hz, 1H), 7.01 (d, *J* = 7.4 Hz, 1H), 5.79 (s, 1H), 5.42 (s, 1H), 2.93 (t, *J* = 7.5 Hz, 2H), 2.79 (t, *J* =7.4 Hz, 2H), 7.12-2.06 (m, 2H), 2.04 (s, 3H).

**$^{13}$C NMR (100 MHz, CDCl$_3$)**: δ 166.3, 145.1, 140.6, 134.5, 133.7, 127.0, 120.7, 119.8, 118.9, 33.1, 29.9, 24.7, 18.6.
**HRMS (+ESI)**: Calculated: 202.1226 ($C_{13}H_{16}NO$). Observed: 202.1224.

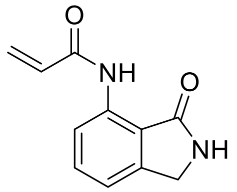

***N*-(3-oxoisoindolin-4-yl)acrylamide (TRH 1-152).**
To a solution of 7-aminoisoindolin-1-one (99 mg, 0.67 mmol) in dichloromethane (4 mL) was added acryloyl chloride (0.07 mL, 0.8 mmol) followed by triethylamine (0.11 mL, 0.8 mmol) at 0˚ C under $N_2$ atmosphere. After stirring for 20 minutes, the reaction mixture was warmed to room temperature and stirred overnight. The reaction mixture was washed twice with brine, dried over magnesium sulfate, and concentrated *in vacuo*. The resulting crude material was purified by silica gel chromatography (50 to 60% ethyl acetate in hexanes) to afford the title compound (58 mg, 43%) as a white solid.
**$^1$H NMR (400 MHz, CDCl$_3$)**: δ 10.50 (s, 1H), 8.58 (d, *J* = 8.2 Hz, 1H), 7.55 (t, *J* = 7.9 Hz, 1H), 7.15 (d, *J* = 7.5 Hz, 1H), 6.82 (s, 1H), 6.46 (dd, *J* = 1.3, 17.0 Hz, 1H), 6.36 (dd, *J* = 10.0, 17.0 Hz, 1H), 5.81 (dd, *J* = 1.3, 10.0 Hz, 1H), 4.46 (s, 2H).
**$^{13}$C NMR (100 MHz, CDCl$_3$)**: δ 172.9, 164.2, 143.9, 138.2, 133.8, 131.8, 127.8, 118.0, 117.7, 117.6, 45.6.
**HRMS (+ESI)**: Calculated: 203.0815 ($C_{11}H_{11}N_2O_2$). Observed: 203.0814.

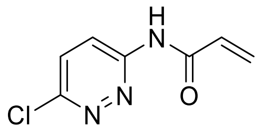

***N*-(6-chloropyridazin-3-yl)acrylamide (TRH 1-155).**
To a solution 3-amino-6-chloropyridazine (261 mg, 2.0 mmol) in dichloromethane (10 mL) was added acryloyl chloride (0.20 mL, 2.4 mmol) followed by triethylamine (0.34 mL, 2.4 mmol) at 0˚C under an atmosphere of nitrogen. After stirring for 20 minutes, the reaction mixture was warmed to room temperature and stirred overnight. The solution was washed twice with brine, dried over magnesium sulfate, and concentrated *in vacuo*. The resulting crude material was purified by silica gel chromatography (40% to 50% ethyl acetate in hexanes) to yield the product (23 mg, 6%) as a pale-yellow solid.
**$^1$H NMR (400 MHz, CDCl$_3$)**: δ 10.06 (s, 1H), 8.70 (d, *J* = 9.4 Hz, 1H), 7.57 (d, *J* = 9.4 Hz, 1H), 6.73 (dd, *J* = 10.2, 16.8 Hz, 1H) 6.56 (dd, *J* = 1.2, 16.8, 1H), 5.94 (dd, *J* = 1.2, 10.2 Hz, 1H).
**$^{13}$C NMR (100 MHz, CDCl$_3$)**: δ 164.8, 155.2, 152.3, 130.7, 130.4, 130.3, 122.0.
**HRMS (+ESI)**: Calculated: 182.0127 ($C_7H_5N_3OCl$). Observed: 182.0126

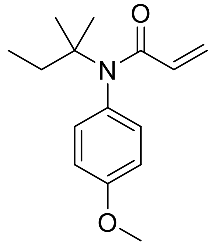

***N*-(4-methoxyphenyl)-*N*-(*tert*-pentyl)acrylamide (TRH 1-170).**
To a solution of 4-methoxy-*N*-(*tert*-pentyl)aniline (94 mg, 0.49 mmol) in dichloromethane (5 mL) was added acryloyl chloride (0.05 mL, 0.6 mmol) followed by triethylamine (0.09 mL, 0.6 mmol) at 0˚C under an atmosphere of nitrogen. After stirring for 15 minutes, the reaction mixture was warmed to room temperature and stirred an additional 18 hours. The reaction mixture was then washed with saturated aqueous sodium bicarbonate solution, brine, dried over magnesium sulfate, and concentrated *in vacuo*. The resulting crude material was purified by silica gel chromatography (0% to 20% ethyl acetate in hexanes) to yield the title compound (82 mg, 68%) as a pale-yellow oil.

**$^1$H NMR (400 MHz, $CDCl_3$)**: δ 6.99 (d, *J* = 8.7 Hz, 2H), 6.85 (d, *J* = 8.7 Hz, 2H), 6.17 (dd, *J* = 1.9, 16.7 Hz, 1H), 5.76 (dd, *J* = 10.3, 16.7 Hz, 1H), 5.28 (dd, *J* = 1.9, 10.3 Hz, 1H), 3.81 (s, 3H), 2.11 (q, *J* = 7.5 Hz, 2H), 1.20 (s, 6H), 0.91 (t, *J* = 7.5 Hz, 3H).
**$^{13}$C NMR (100 MHz, $CDCl_3$)**: δ 166.3, 159.0, 134.3, 131.49, 131.45, 125.6, 114.1, 61.7, 55.5, 32.0, 27.4, 9.4.
**HRMS (+ESI)**: Calculated: 247.1572 ($C_{15}H_{21}NO_2$). Observed: 247.1577.

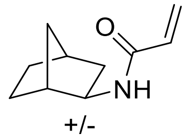


***N*-(*exo*-norborn-2-yl)acrylamide (TRH 1-176).**
To a solution of *exo*-2-aminonorbornane (0.24 mL, 2 mmol) in dichloromethane (10 mL) was added acryloyl chloride (0.20 mL, 2.4 mmol) followed by triethylamine (0.33 mL, 2.4 mmol) at 0˚C under an atmosphere of nitrogen. After stirring for 20 minutes, the reaction mixture was warmed to room temperature and stirred for an additional 18 hours. The reaction mixture was then washed with saturated aqueous sodium bicarbonate solution, brine, dried over magnesium sulfate, and concentrated *in vacuo*. The resulting crude material was purified by silica gel chromatography (30% ethyl acetate in hexanes) to yield the title compound (271 mg, 82%) as a white solid.
**$^1$H NMR (400 MHz, $CDCl_3$)**: δ 6.42 (s, 1H), 6.25 (dd, *J* = 2.3, 17.0 Hz, 1H), 6.18 (dd, *J* = 9.5, 17.0 Hz, 1H), 5.58 (dd, *J* = 2.3, 9.5 Hz, 1H), 3.8-3.77 (m, 1H), 2.27-2.24 (m, 2H), 1.78 (ddd, *J* = 2.1, 8.1, 13.0 Hz, 1H), 1.55-1.38 (m, 3H), 1.30-1.10 (m, 4H).
**$^{13}$C NMR (100 MHz, $CDCl_3$)**: δ 165.0, 131.4, 125.8, 52.9, 42.4, 40.0, 35.7, 35.6, 28.2, 26.6.
**HRMS (+EI)**: Calculated:165.1154 ($C_{10}H_{15}NO$). Observed: 165.1155.

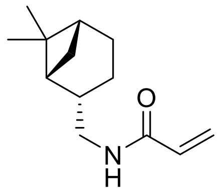


***N*-(((1*R*,2*S*,5*R*)-6,6-dimethylbicyclo[3.1.1]heptan-2-yl)methyl)acrylamide (TRH 1-178).**
To a solution of (-)-*cis*-myrtanylamine (0.34 mL, 2 mmol) in dichloromethane (10 mL) was added acryloyl chloride (0.20 mL, 2.4 mmol) followed by triethylamine (0.33 mL, 2.4 mmol) at 0˚C under an atmosphere of nitrogen. After stirring for 20 minutes, the reaction mixture was warmed to room temperature and stirred for an additional 21 hours. The reaction mixture was then washed with saturated aqueous sodium bicarbonate solution, brine, dried over magnesium sulfate, and concentrated *in vacuo*. The resulting crude material was purified by silica gel chromatography (20 to 30% ethyl acetate in hexanes) to yield the title compound (369 mg, 89%) as a white solid.
**$^1$H NMR (600 MHz, $CDCl_3$)**: δ 6.26 (dd, *J* = 1.5, 17.0 Hz, 1H), 6.11 (dd, *J* = 10.3, 17.0 Hz, 1H) 5.85 (s, 1H), 5.61 (dd, *J* = 1.5, 10.3 Hz, 1H), 3.39-3.29 (m, 2H), 2.38-2.34 (m, 1H), 2.26-2.21 (m, 1H), 1.98-1.90 (m, 4H), 1.88-1.83 (m, 1H), 1.53-1.47 (m, 1H), 1.19 (s, 3H), 1.04 (s, 3H), 0.89 (d, *J* = 9.6 Hz, 1H).
**$^{13}$C NMR (150 MHz, $CDCl_3$)**: δ 165.7, 131.2, 126.2, 45.3, 43.9, 41.5, 38.8, 33.3, 28.1, 26.1, 23.3, 19.9.
**HRMS (-ESI)**: Calculated: 206.1550 ($C_{13}H_{20}NO$). Observed: 206.1551.

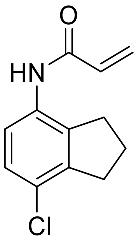


**N-(7-chloro-2,3-dihydro-1H-inden-4-yl)acrylamide (YP 1-1)**
A solution of *N*-(2,3-dihydro-1H-inden-4-yl)acrylamide (187 mg, 1.0 mmol) in PEG 400 (5.2 mL) was cooled to 0˚C and *N*-chlorosuccinimide (140 mg, 1.0 mmol) added. After 30 minutes, the mixture was warmed to room temperature and stirred overnight. The reaction mixture was diluted with ethyl acetate, washed with brine twice, and dried over magnesium sulfate. The volatiles were removed *in vacuo* and the crude product purified by silica

gel chromatography (30% ethyl acetate in hexanes). The obtained mixture of isomers were recrystallized to afford the title compound (47 mg, 22% yield) as a white solid.

**$^{1}$H NMR (400MHz, $CDCl_3$)**: δ 7.78 (d, *J* = 8.8 Hz, 1H), 7.15-7.11 (m, 2H), 6.42 (dd, *J* = 1.4, 16.8 Hz, 1H), 6.26 (dd, *J* = 10.2, 16.8 Hz, 1H), 5.77 (dd, *J* = 1.4, 10.2 Hz, 1H), 2.98 (t, *J* = 7.6 Hz, 2H), 2.87 (t, *J* = 7.5 Hz, 2H), 2.12 (quint, *J* = 7.5 Hz, 2 H).

**$^{13}$C NMR (100MHz, $CDCl_3$)**: δ 163.4, 143.1, 136.1, 132.2, 131.0, 128.0, 127.2, 126.7, 120.9, 32.7, 31.1, 24.0.

**HRMS (+ESI)**: Calculated: 220.0535 ($C_{12}H_{11}ClNO$). Observed: 220.0533.

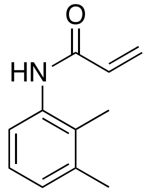

**N-(2,3-dimethylphenyl)acrylamide (YP 1-18)**

A solution of 2,3-dimethylaniline (121 mg, 1.0 mmol) in DCM (10 mL) was cooled to 0°C and acryloyl chloride (109 mg, 1.2 mmol) and triethylamine (121 mg, 1.2 mmol) were added sequentially. The reaction mixture was maintained at this temperature for 30 minutes and then warmed to room temperature and stirred overnight. The reaction mixture was washed twice with brine and dried over magnesium sulfate. Volatiles were removed *in vacuo* and the crude material purified by silica gel chromatography (30% to 40% ethyl acetate in hexanes) to afford the product (154 mg, 88%) as a white solid.

**$^{1}$H NMR (400MHz, $CDCl_3$)**: δ 7.49 (d, *J* = 7.9 Hz, 1H), 7.29 (s, 1H), 7.11-7.07 (m, 1H), 7.01 (d, *J* = 7.7, 1H) 6.40 (d, *J* = 17.1, 1H), 6.30 (dd, *J* = 7.3, 17.1 Hz, 1H), 5.74 (d, *J* = 10.1 Hz, 1H), 2.29 (s, 1H), 2.13 (s, 1H).

**$^{13}$C NMR (100MHz, $CDCl_3$)**: δ 135.1, 131.2, 127.6, 127.3, 125.9, 122.3, 20.6, 13.9.

**HRMS (+ESI)**: Calculated: 176.1070 ($C_{11}H_{14}NO$). Observed: 176.1068.

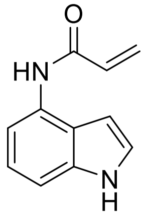

**N-(1H-indol-4-yl)acrylamide (YP 1-19)**

A solution of 4-aminoindole (132 mg, 1 mmol) in DCM/DMF (1:1 *v:v*, 10 mL) was cooled to 0°C and acryloyl chloride (109 mg, 1.2 mmol) and triethylamine (121 mg, 1.2 mmol) added sequentially. The reaction mixture was stirred at this temperature for 26 minutes and then warmed to room temperature and stirred overnight. The reaction mixture was washed twice with brine and dried over magnesium sulfate. Volatiles were removed *in vacuo* and the crude product purified by basic alumina chromatography (60% to 75% ethyl acetate in hexanes) to afford the title compound (56mg, 30%) as a white-grey solid.

**$^{1}$H NMR (600MHz, MeOD)**: δ 7.51 (d, *J* = 7.6 Hz, 1H), 7.24-7.22 (m, 2H), 7.08 (t, *J* = 7.6 Hz, 1H), 6.64 (dd, *J* = 10.1, 16.7 Hz, 2H), 6.38 (dd, *J* = 1.7, 16.9 Hz, 1H), 5.78 (dd, *J* = 1.7, 10.3 Hz, 1H), 4.6 (s, 1H).

**$^{13}$C NMR (150MHz, MeOD)**: δ 165.0, 137.2, 131.1, 129.2, 126.0, 123.8, 121.5, 120.9, 112.2, 108.4, 98.5.

**HRMS (+ESI)**: Calculated: 187.0866 ($C_{11}H_{11}N_2O$). Observed: 187.0865.

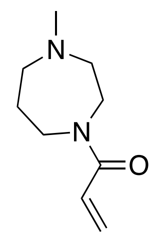

**1-(4-methyl-1,4-diazepan-1-yl)prop-2-en-1-one (YP 1-23)**, mixture of rotamers
A solution of 1-methylhomopiperazine (114 mg, 1.0 mmol) in DCM (10 mL) was cooled to 0 °C and acryloyl chloride (109 mg, 1.2 mmol) and triethylamine (121 mg, 1.2 mmol) added sequentially. The solution was maintained at this temperature for 30 minutes and then warmed to room temperature and stirred overnight. The reaction mixture was washed twice with brine and dried over magnesium sulfate. After removal of the volatiles in *vacuo*, the crude product was purified via silica gel chromatography (1% to 10% methanol in DCM) affording the title compound (58 mg, 51%) as a yellow oil.
**$^1$H NMR (400MHz, $CDCl_3$):** δ 6.61-6.53 (m, 1H), 6.35-6.29 (m, 1H), 5.70-5.66 (m, 1H), 3.74-3.72 (m, 1 H), 3.69 (t, *J* = 6.4 Hz, 1H), 3.65-3.61 (m, 2H), 2.66-2.63 (m, 2H), 2.59-2.54 (m, 2H), 2.37 (s, 3H), 1.94 (quint, *J* = 6.2 Hz, 2H).
**$^{13}$C NMR (100MHz, $CDCl_3$)**: δ 166.4, 166.3, 128.0, 127.9, 127.8, 127.6, 59.1, 58.0, 57.1, 56.8, 47.4, 47.1, 46.7, 46.6, 45.3, 44.8, 28.1, 26.9.
**HRMS (+ESI)**: Calculated: 169.1335 ($C_9H_{17}N_2O$). Observed: 169.1333.

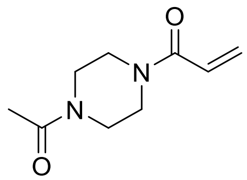

**1-(4-acetylpiperazin-1-yl)prop-2-en-1-one (YP 1-24)**
A solution of 1-acetylpiperazine (128 mg, 1.0 mmol) in DCM (10 mL) was cooled to 0 °C and acryloyl chloride (109 mg, 1.2 mmol) and triethylamine (121 mg, 1.2 mmol) added sequentially. The solution was stirred at this temperature for 23 minutes and then warmed to room temperature and stirred an additional two hours. The reaction mixture was washed twice with brine, dried over magnesium sulfate, and the volatiles removed *in vacuo*. The crude material was purified via silica gel chromatography (0% to 10% methanol in DCM) to afford the title compound (40 mg, 18%) as a yellow oil.
**$^1$H NMR (400MHz, $CDCl_3$)**: δ 6.57 (dd, *J* = 10.5, 16.8 Hz, 1H), 6.33 (dd, *J* = 1.8, 16.8 Hz, 1H), 5.75 (dd, *J* = 1.9, 10.5 Hz, 1H), 3.72 (s, 1H), 3.66-3.64 (m, 3H), 3.57 (s, 1H), 3.51-3.49 (m, 2H), 2.13 (s, 3H).
**$^{13}$C NMR (100MHz, $CDCl_3$)**: δ 169.0, 165.6, 128.7, 127.0, 41.9, 41.4, 21.4.
**HRMS (+ESI)**: Calculated: 183.1128 ($C_9H_{15}N_2O_2$). Observed: 183.1126.

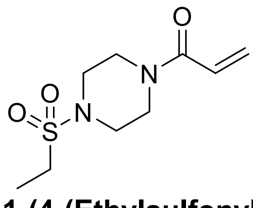

**1-(4-(Ethylsulfonyl)piperazin-1-yl)prop-2-en-1-one (YP 1-25)**
A solution of 1-(ethanesulfonyl)piperazine (178 mg, 1.0 mmol) in DCM (10 mL) was cooled to 0 °C and acryloyl chloride (109 mg, 1.2 mmol) and triethylamine (121 mg, 1.2 mmol) added sequentially. The solution was stirred at this temperature for 27 minutes and then warmed to room temperature and stirred for an additional two hours. The solution was washed twice with brine and dried over magnesium sulfate. The crude material was purified by silica gel chromatography (1% to 10% methanol in DCM) to afford the title compound (163 mg, 70%) as a white-yellow solid.
**$^1$H NMR (400MHz, $CDCl_3$)**: δ 6.57 (dd, *J* = 10.5, 16.8 Hz, 1H), 6.32 (dd, *J* = 1.9, 16.8 Hz, 1H), 5.76 (dd, *J* = 1.8, 10.5 Hz, 1H), 3.77 (s, 2H), 3.67 (s, 2H), 3.32 (t, *J* = 5.2 Hz, 4H), 2.98 (q, *J* = 7.5 Hz, 2H), 1.37 (t, *J* = 7.4, 3H).
**$^{13}$C NMR (100MHz, $CDCl_3$)**: δ 165.5, 128.8, 127.0, 77.4, 45.9, 45.6, 44.2, 41.9, 7.8.
**HRMS (+ESI)**: Calculated: 233.0954 ($C_9H_{17}N_2O_3S_1$). Observed: 233.0953.

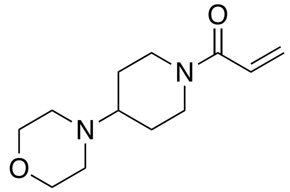

**1-(4-morpholinopiperidin-1-yl)prop-2-en-1-one (YP 1-42)**
Following **General Procedure A** using 4-morpholinopiperidine (336 mg, 2.0 mmol), the product was obtained after silica gel chromatography (1% methanol and 80% ethyl acetate in hexanes) as a colorless oil (259 mg, 58%).
**$^{1}$H NMR (400MHz, $CDCl_3$)**: δ 6.42 (dd, *J* = 10.6, 16.8 Hz, 1H), 6.06 (dd, *J* = 2.0, 16.8 Hz, 1H), 5.49 (dd, *J* = 2.0, 10.6 Hz, 1H), 4.45 (d, *J* = 12.8 Hz, 1H), 3.86 (d, *J* = 12.8 Hz, 1H), 3.52 (t, *J* = 4.7 Hz, 4H), 2.90 (t, *J* = 12.8 Hz, 1H), 2.55-2.48 (m, 1H), 2.37-2.35 (m, 4H), 2.26 (tt, *J* = 3.7, 11.0 Hz, 1H), 1.72 (d, *J* = 12.8 Hz, 2H), 1.30-1.20 (m, 2H).
**$^{13}$C NMR (100MHz, $CDCl_3$)**: δ 165.0, 127.7, 127.3, 67.1, 61.6, 49.6, 44.9, 41.1, 28.9, 27.8.
**HRMS (+ESI)**: Calculated: 225.1598 ($C_{12}H_{21}N_2O_2$). Observed: 225.1595.

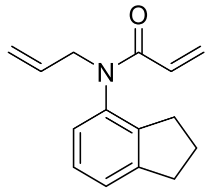

**N-allyl-N-(2,3-dihydro-1H-inden-4-yl)acrylamide (IGA 1-12)**
To a solution of sodium hydride (96 mg, 4.0 mmol) in tetrahydrofuran (8 mL) was added *N*-(2,3-dihydro-1H-inden-4-yl)acrylamide (187 mg, 1.0 mmol) in tetrahydrofuran (2 mL) under an atmosphere of nitrogen. The reaction mixture was cooled to 0°C and 3-bromoprop-1-ene (484 mg, 4.0 mmol) was and the mixture warmed to room temperature and stirred overnight. The reaction was quenched by the addition of water and extracted with ethyl acetate. The crude product was purified by silica gel chromatography (20% ethyl acetate in hexanes) to afford the product (151 mg, 67%) as a yellow crystalline solid.
**$^{1}$H NMR (400MHz, $CDCl_3$)**: δ 7.06-7.18 (m, 2H), 6.80-6.88 (m, 1H), 6.26-6.37 (dd, *J* = 16.8, 2.0 Hz, 1H), 5.76-5.96 (m, 2H), 5.38-5.48 (dd, *J* = 10.3, 2.1 Hz, 1H), 4.98-5.08 (m, 2H), 4.40-4.52 (ddt, *J* = 14.5, 6.3, 1.3 Hz, 1H), 4.00-4.11 (ddt, *J* = 14.5, 6.8, 1.2 Hz, 1H), 2.82-2.98 (m, 2H), 2.59-2.79 (m, 2H), 1.92-2.07 (m, 2H).
**$^{13}$C NMR (100MHz, $CDCl_3$)**: δ 165.1, 146.5, 142.4, 137.9, 133.0, 128.4, 127.8, 127.48, 126.1, 124.3, 118.1, 51.6, 33.3, 30.9, 25.0.
**HRMS (+ESI)**: Calculated: 228.13 ($C_{15}H_{17}NO$). Observed: 228.1381.

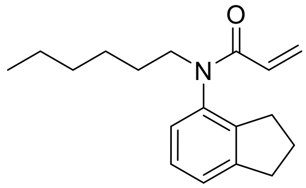

**N-allyl-N-(2,3-dihydro-1H-inden-4-yl)acrylamide (IGA 1-15)**
To a solution of sodium hydride (96 mg, 4.0 mmol) in tetrahydrofuran (8 mL) was added *N*-(2,3-dihydro-1H-inden-4-yl)acrylamide (187 mg, 1.0 mmol) in tetrahydrofuran (2 mL) under an atmosphere of nitrogen. The solution was cooled to 0°C and 1-bromohexane (660 mg, 4.0 mmol) was added after which point the solution was warmed to room temperature and stirred overnight. The solution was quenched with water and extracted with ethyl acetate. The crude product was purified via silica gel chromatography (20% ethyl acetate in hexanes) to afford the product in 34% yield as a yellow oil (92 mg).
**$^{1}$H NMR (400MHz, $CDCl_3$)**: δ 7.11-7.25 (m, 2H), 6.86-6.96 (dd, *J* = 7.5, 1.2 Hz, 1H), 6.30-6.40 (dd, *J* = 16.8, 2.1 Hz, 1H), 5.86-6.00 (m, 1H), 5.41-5.51 (dd, *J* = 10.3, 2.1 Hz, 1H), 3.82-3.96 (m, 1H), 3.42-3.56 (m, 1H), 2.90-3.04 (m, 2H), 2.65-2.85 (m, 2H), 1.98-2.16 (m, 2H), 1.47-1.63 (m, 2H), 1.20-1.36 (m, 6H), 0.80-0.90 (m, 3H).
**$^{13}$C NMR (100MHz, $CDCl_3$)**: δ 165.2, 146.5, 142.4, 138.2, 128.6, 127.5, 127.4, 126.1, 124.1, 48.7, 33.3, 31.6, 30.9, 27.9, 26.7, 25.0, 22.6, 14.1.
**HRMS (+ESI)**: Calculated: 272.19 ($C_{18}H_{25}NO$). Observed: 272.2007.

### Supporting Tables

**Table S1. IsoTOP-ABPP analysis of nimbolide treatment *in situ* in 231MFP breast cancer cells.** IsoTOP-ABPP analysis of nimbolide treatment *in situ* (10 μM). 231MFP breast cancer cells were treated with DMSO or nimbolide (10 μM, 1.5 h *in situ*), after which cells were harvested and proteomes were labeled *ex situ* with IA-alkyne (100 μM, 1 h), followed by appendage of isotopically light (for DMSO-treated) or heavy (for nimbolide-treated) TEV protease cleavable biotin-azide tags by copper-catalyzed azide-alkyne cycloaddition (CuAAC). Control and treated proteomes were subsequently combined in a 1:1 ratio, probe-labeled proteins were avidin-enriched, digested with trypsin, and probe-modified tryptic peptides were eluted by TEV protease, analyzed by LC-MS/MS, and light to heavy probe-modified peptide ratios were quantified. Shown are data from n=3 biological replicates/group.
**Tab 1.** Total isoTOP-ABPP proteomic dataset
**Tab 2.** Analyzed isoTOP-ABPP dataset. For those probe-modified peptides that showed ratios >2, we only interpreted those targets that were present across all three biological replicates, were statistically significant, and showed good quality MS1 peak shapes across all biological replicates. Light versus heavy isotopic probe-modified peptide ratios are calculated by taking the mean of the ratios of each replicate paired light vs. heavy precursor abundance for all peptide spectral matches (PSM) associated with a peptide. The paired abundances were also used to calculate a paired sample t-test p-value in an effort to estimate constancy within paired abundances and significance in change between treatment and control. P-values were corrected using the Benjamini/Hochberg method.

**Table S2. TMT-based quantitative proteomic analysis of proteins enriched by nimbolide-alkyne probe *in situ* treatment in 231MFP breast cancer cells.** 231MFP breast cancer cells were treated with DMSO vehicle or the nimbolide-alkyne probe (50 μM) for 1.5 h. Probe-labeled proteins were conjugated to biotin-azide by CuAAC and subsequently avidin-enriched from 231MFP proteomes and tryptic digests from enriched proteins were analyzed by TMT-based quantitative proteomics. Shown are the proteins from this experiment that showed at least 2 unique peptides, as well as TMT ratios of no-probe versus probe. The data shown are from n=3 biological replicates/group.

**Table S3. TMT-based quantitative proteomic profiling of XH2-treatment in 231MFP breast cancer cells.** Tandem mass tag (TMT)-based quantitative proteomic profiling of 231MFP breast cancer cells treated with DMSO vehicle or nimbolide (100 nM) **(Tab 1)** or XH2 (100 nM) **(Tab 2)** for 12 h. Shown are the data for proteins identified that showed at least 2 unique peptides from n=3 biological replicates/group.

**Table S4. Structures of covalent ligands screened against RNF114**

**Supporting Figures**

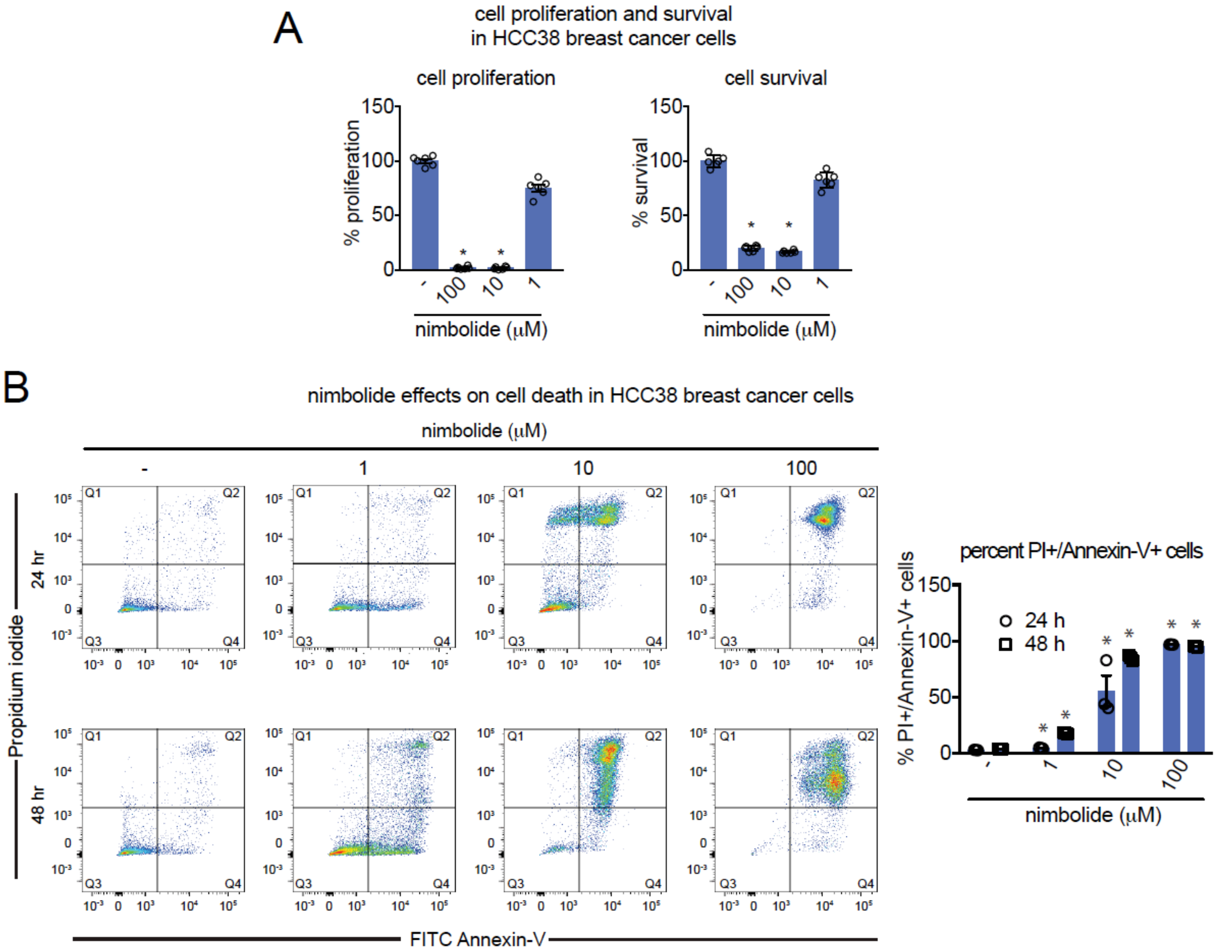


**Figure S1. Nimbolide impairs HCC38 breast cancer cell proliferation or survival. (A)** HCC38 breast cancer cell proliferation in serum-containing media and serum-free cell survival. Cells were treated with DMSO vehicle or nimbolide and cell viability was assessed after 48 h by Hoechst stain. **(B)** Percent of propidium iodide and Annexin-V-positive (PI+/Annexin-V+) cells assessed by flow cytometry after treating HCC38 cells with DMSO vehicle or nimbolide for 24 or 48 h. Shown on the left panels are representative FACS data. On the right bar graph are noted late-stage apoptotic cells defined as defined as PI+/Annexin-V+ cells. Data shown in **(A** and **B)** are average ± sem, n=3-6 biological replicates/group. Statistical significance was calculated with unpaired two-tailed Student's t-tests. Significance is expressed as *p<0.05 compared to vehicle-treated controls.

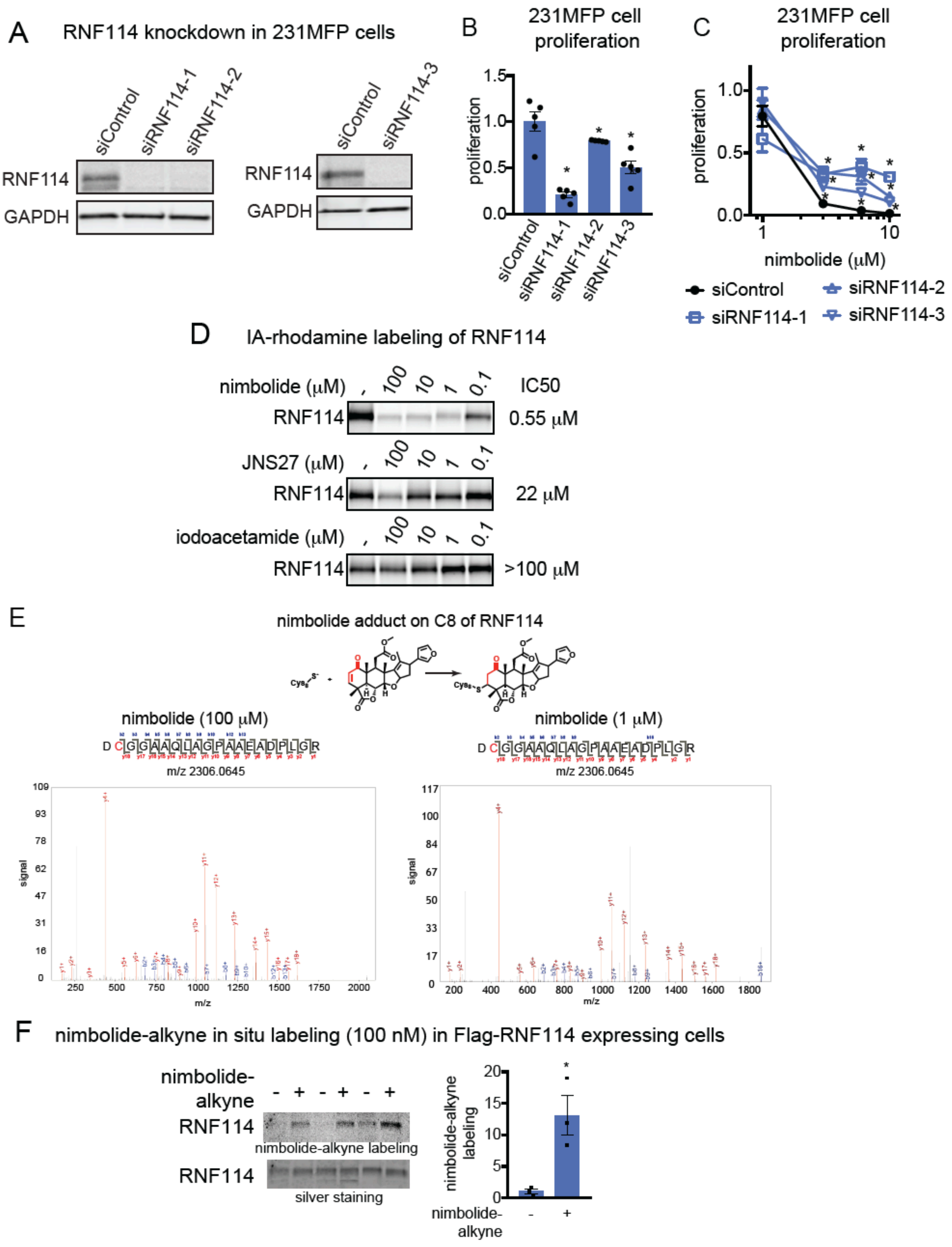


**Figure S2. Elucidating the Role of RNF114 in nimbolide-mediated effects. (A)** RNF114 knockdown by 3 independent siRNAs targeting RNF114 validated by Western blotting of RNF114 compared to siControl 231MFP cells. GAPDH expression is shown as a loading control. **(B)** 231MFP cell proliferation after 24 h in siControl and siRNF114 cells assessed by Hoechst stain. **(C)** Nimbolide effects on 231MFP siControl and siRNF114 231MFP breast cancer cells. Cells were treated with DMSO vehicle or nimbolide for 24 h after which proliferation was assessed by Hoechst stain. Data for each siControl or siRNF114 group was normalized to the respective DMSO vehicle control in each group. **(D)** Gel-based ABPP analysis of nimbolide, JNS27, and iodoacetamide competition against IA-rhodamine labeling of recombinant human RNF114 protein. RNF114 protein was pre-treated with DMSO vehicle or nimbolide, JNS27, or iodoacetamide for 30 min prior to labeling of RNF114 with IA-rhodamine (100 nM) for 30 min. RNF114 IA-rhodamine labeling was assessed by SDS/PAGE and in-gel fluorescence. **(E)** Pure RNF114 protein was labeled with nimbolide (100 or 1 μM, 1 h) and subjected to tryptic digestion and LC-MS/MS analysis. Shown is the nimbolide-modified adduct on C8 of RNF114. **(F)** Nimbolide-alkyne *in situ* labeling. 231MFP cells stably expressing a Flag-tagged RNF114 were treated with DMSO vehicle or nimbolide-alkyne (100 nM) for 4 h. RNF114 was subsequently enriched from harvested cell lysates and then rhodamine-azide was appended onto probe-labeled proteins by CuAAC, after which nimbolide-alkyne labeling was visualized by SDS/PAGE and in-gel fluorescence. RNF114 enrichment and loading were assessed by silver staining. RNF114 labeling by the nimbolide-alkyne probe were quantified by densitometry. Gels shown in **(A** and **D)** are representative gels from n=3/group. Data shown in **(B** and **C)** are average ± sem, n=3-5 biological replicates/group. Statistical significance was calculated with unpaired two-tailed Student's t-tests. Significance is expressed in **(B)** as *$p<0.05$ compared to siControl cells. Significance in **(C)** is expressed as *$p<0.05$ compared to corresponding concentration of nimbolide treatment in siControl cells. Significance in **(F)** is expressed as *$p<0.05$ compared to vehicle-treated controls.

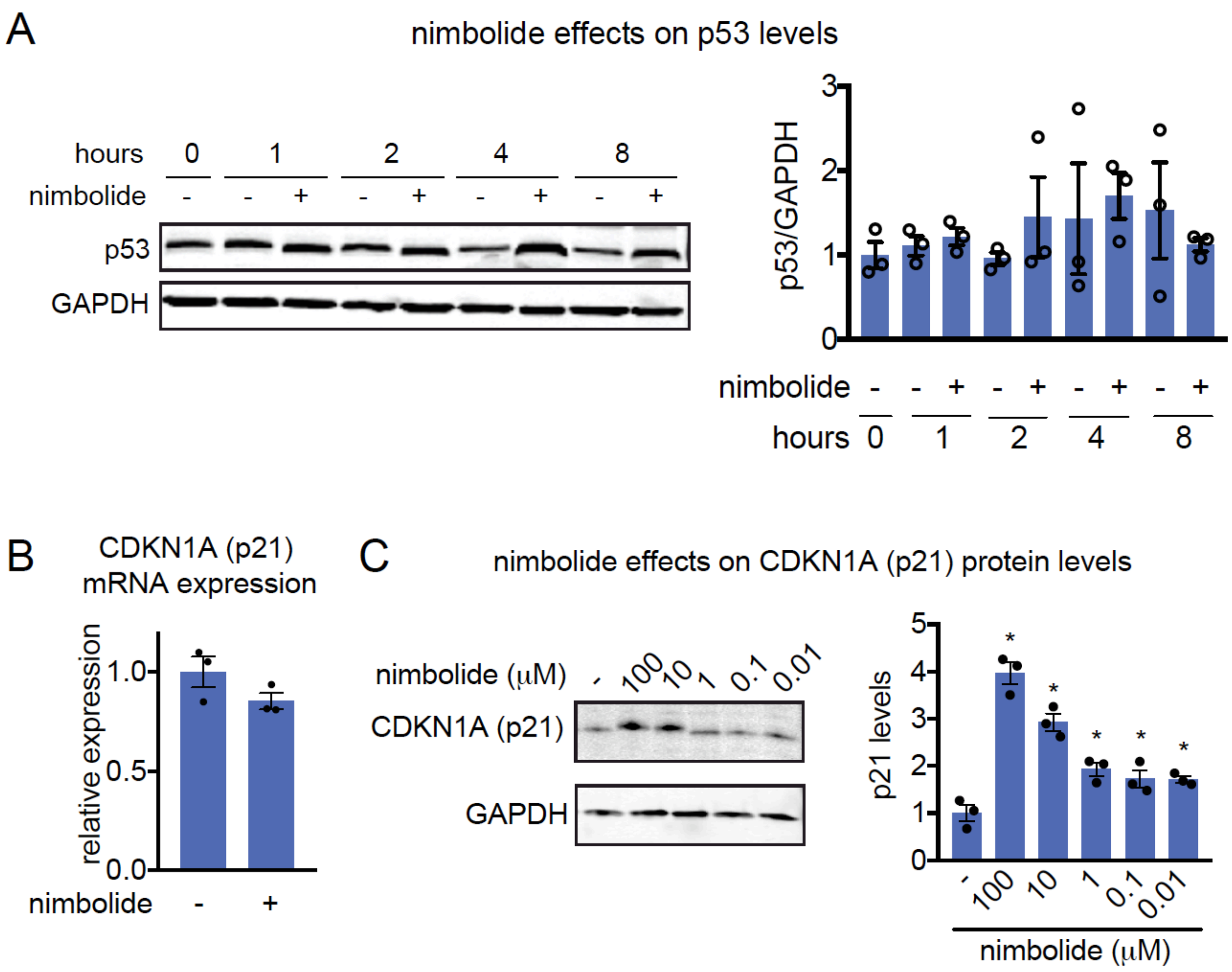


**Figure S3. Levels of p53 and p21 in nimbolide-treated 231MFP breast cancer cells.** **(A)** p53 levels in 231MFP breast cancer cells treated with DMSO vehicle or nimbolide (100 μM) assessed by Western blotting alongside GAPDH as a loading control. **(B)** p21 mRNA expression levels in 231MFP cells treated with nimbolide (100 μM) for 1 h assessed by qPCR. **(C)** Dose-response of p21 elevation with DMSO vehicle or nimbolide treatment for 1 h, assessed by Western blotting alongside GAPDH as a loading control. Gels shown in **(A** and **C)** are representative of an n=3 biological replicates/group. Blots were quantified by densitometry and normalized to loading control. Data shown in bar graph are average ± sem. Significance expressed in **(C)** as *p<0.05 compared to vehicle-treated controls.

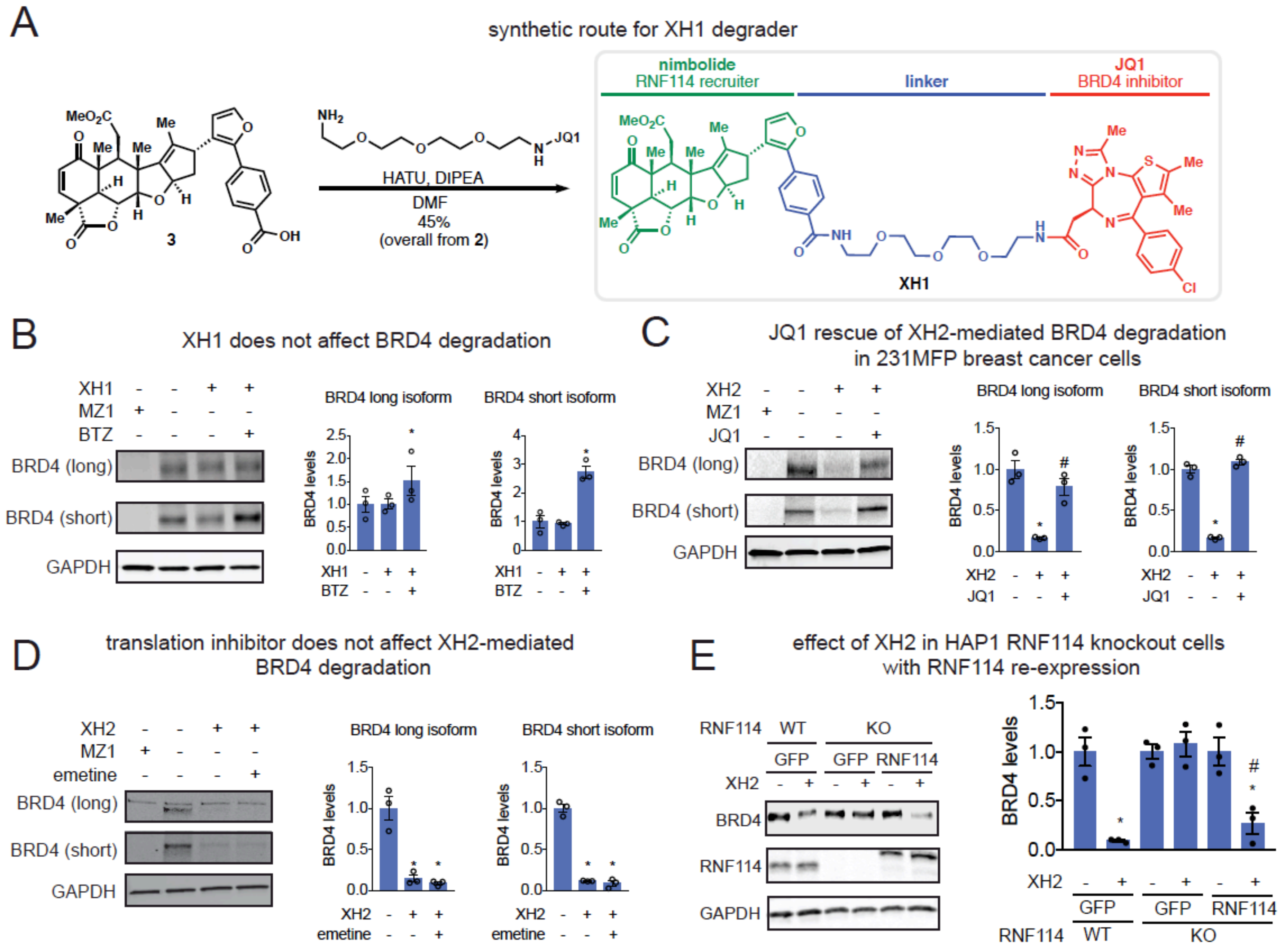


**Figure S4. Characterization of nimbolide-based degraders. (A)** Synthetic route for XH1 degrader. **(B)** BRD4 levels in 231MFP breast cancer cells pre-treated with DMSO vehicle or proteasome inhibitor bortezomib (1 μM) for 30 min prior to and also during XH1 treatment (100 nM) for 12 h, assessed by Western blotting alongside GAPDH loading control. **(C-D)** BRD4 levels in 231MFP breast cancer cells pre-treated with DMSO vehicle, JQ1 (1 μM) **(C)**, or emetine (75 μM) **(D)** for 30 min prior to and also during XH2 treatment (100 nM) for 12 h, assessed by Western blotting alongside a GAPDH loading control. **(E)** Effect of DMSO vehicle or XH2 (100 nM) treatment for 12 h on BRD4 expression in HAP1 RNF114 wild-type and knockout cells transiently transfected with GFP control or wild-type Flag-tagged RNF114. BRD4, RNF114, and GAPDH expression were assessed by Western blotting. Data were normalized to DMSO vehicle controls for each group. Gels shown in **(B-E)** are representative images from n=3/group. Data shown in **(B-E)** are average ± sem. Data in **(B-E)** are n=3 biological replicates/group. Statistical significance was calculated with unpaired two-tailed Student's t-tests. Significance in **(B-D)** is expressed as *p<0.05 compared to the vehicle-treated control groups and #p<0.05 compared to XH1 or XH2-treated groups. Significance in **(E)** is expressed as *p<0.05 compared to vehicle-treated control for each group.

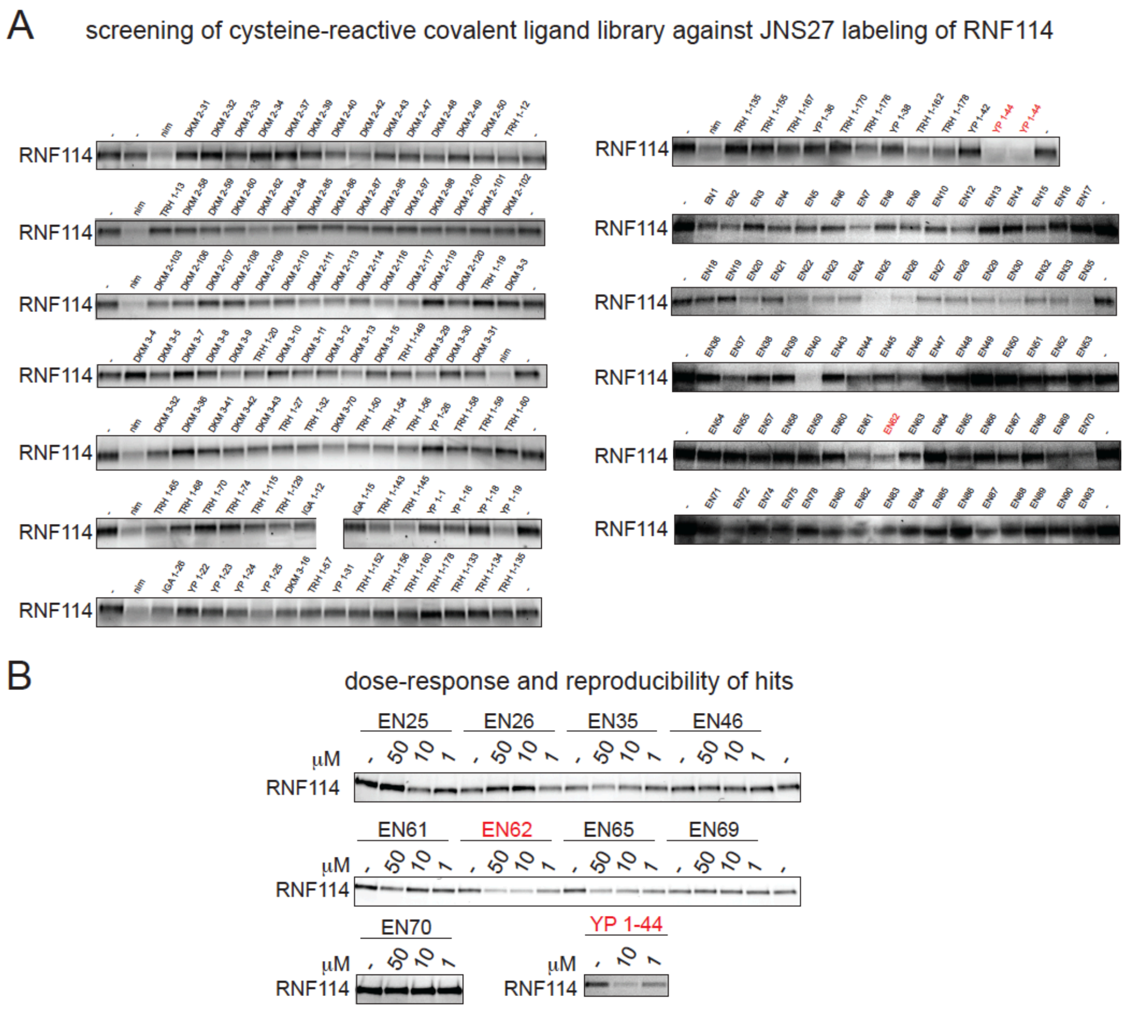


**Figure S5. Gel-based ABPP screen of cysteine-reactive ligands against RNF114. (A)** Cysteine-reactive covalent ligands were screened against JNS27 labeling of pure human RNF114 protein. Covalent ligands (50 µM) were pre-incubated with RNF114 for 30 min prior to labeling with JNS27 for 1 h. Rhodamine-azide was then appended to probe-labeled proteins by CuAAC. Proteins were separated by SDS/PAGE and visualized by in-gel fluorescence. **(B)** Those proteins that showed inhibition of probe-labeling were re-tested in dose-response studies below to identify hits that reproducibly inhibited JNS27 labeling of RNF114 under the same incubation conditions described in **(A)**. Data shown in **(A)** were from n=1. Data shown in **(B)** are representative gels shown from n=3 biological replicates/group.

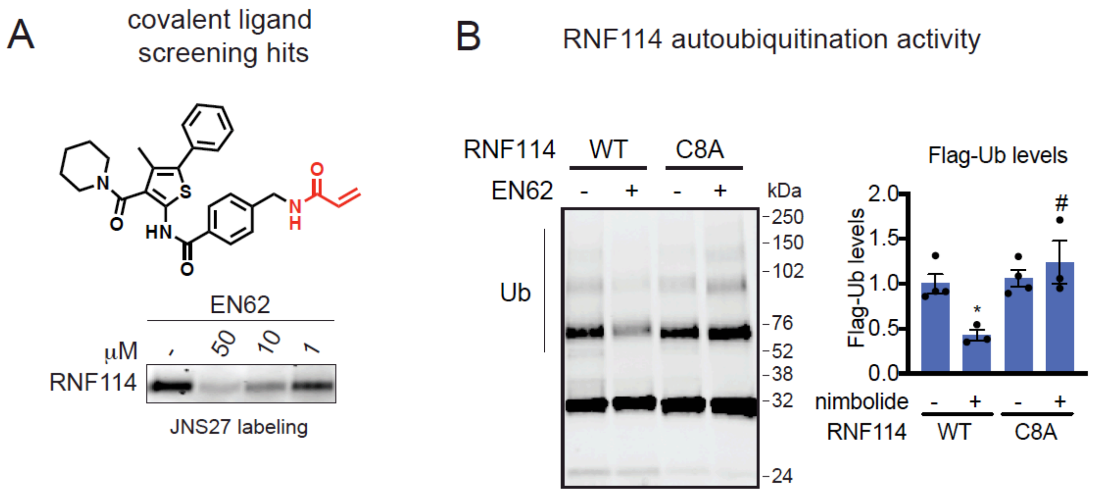


**Figure S6. Chemoproteomics-enabled covalent ligand screening to identify more synthetically tractable covalent ligands against RNF114. (A)** Upon screening a library of cysteine-reactive covalent ligands against JNS27 labeling of RNF114, EN62 was one of the top hits. Shown is the structure of EN62 with the acrylamide reactive moiety highlighted in red. Shown also is a gel-based ABPP analysis of EN62 against JNS27 labeling of pure RNF114. **(B)** RNF114 autoubiquitination assay with DMSO or EN62 (50 μM) treatment with wild-type or C8A mutant RNF114. Gels shown in **(A-B)** are representative images from n=3 biological replicates/group. Significance in **(B)** is expressed as *p<0.05 compared to the vehicle-treated controls. Significance is expressed as #p<0.05 compared to EN62-treated WT RNF114 protein in **(B).**
